# Integrated Interactomics Reveals Novel Protein Associations: The FOXA1-PBX1 Complex as a Case Study

**DOI:** 10.64898/2026.07.16.738938

**Authors:** Zhi Chen, Istvan Szepesi-Nagy, Qingyue Zhang, Srushti Kittane, Yangzhenyu Gao, Juliana Ortiz-Pacheco, Ethan Lane, Justin Hatcher, Jeffrey Estrada, Manor Askenazi, Lan Huang, Beatrix Ueberheide, Ning Zheng, Eneda Toska, Gergely Rona, Michele Pagano

## Abstract

Protein-protein interactions (PPIs) are fundamental to cellular signaling networks, yet many remain undetected due to technical limitations of individual affinity purification approaches. To address this, we systematically mapped the interaction landscapes of six regulatory proteins involved in cell proliferation, immunity, and inflammation, including three transcription factors (TFs) and three kinases. We implemented an integrated proteomics workflow that combined four complementary affinity purification strategies: native immunoprecipitation, two crosslinking-assisted capture methods, and proximity labeling. Combining these approaches revealed distinct yet overlapping interaction profiles, uncovered numerous previously unreported interactors not reliably detected by individual methods, and robustly recovered known interactions while substantially extending PPI networks. Despite method-specific differences at the protein level, functional enrichment analyses showed strong convergence on coherent biological pathways. Biochemical approaches validated most of the previously unreported interactions, including putative weak and transient complexes stabilized by crosslinking. Functional assays revealed a previously unrecognized physical interaction between FOXA1 and PBX1 TFs and demonstrated their cooperative regulation of transcriptional programs and cell fitness in estrogen receptor (ER) positive breast cells. We propose that the FOXA1-PBX1 complex could represent a higher-order regulatory node integrating chromatin accessibility and ER-driven transcriptional output.

**HIGHLIGHTS:**

- Complementary affinity purification strategies uncover putative weak and transient protein-protein interactions
- Functional pathway convergence validates biologically coherent interactome expansion
- Biochemical validations confirm unreported interactions
- Functional validation studies identify a FOXA1-PBX1 pioneer factor complex that regulates estrogen receptor transcriptional programs

**Graphical Abstract:** 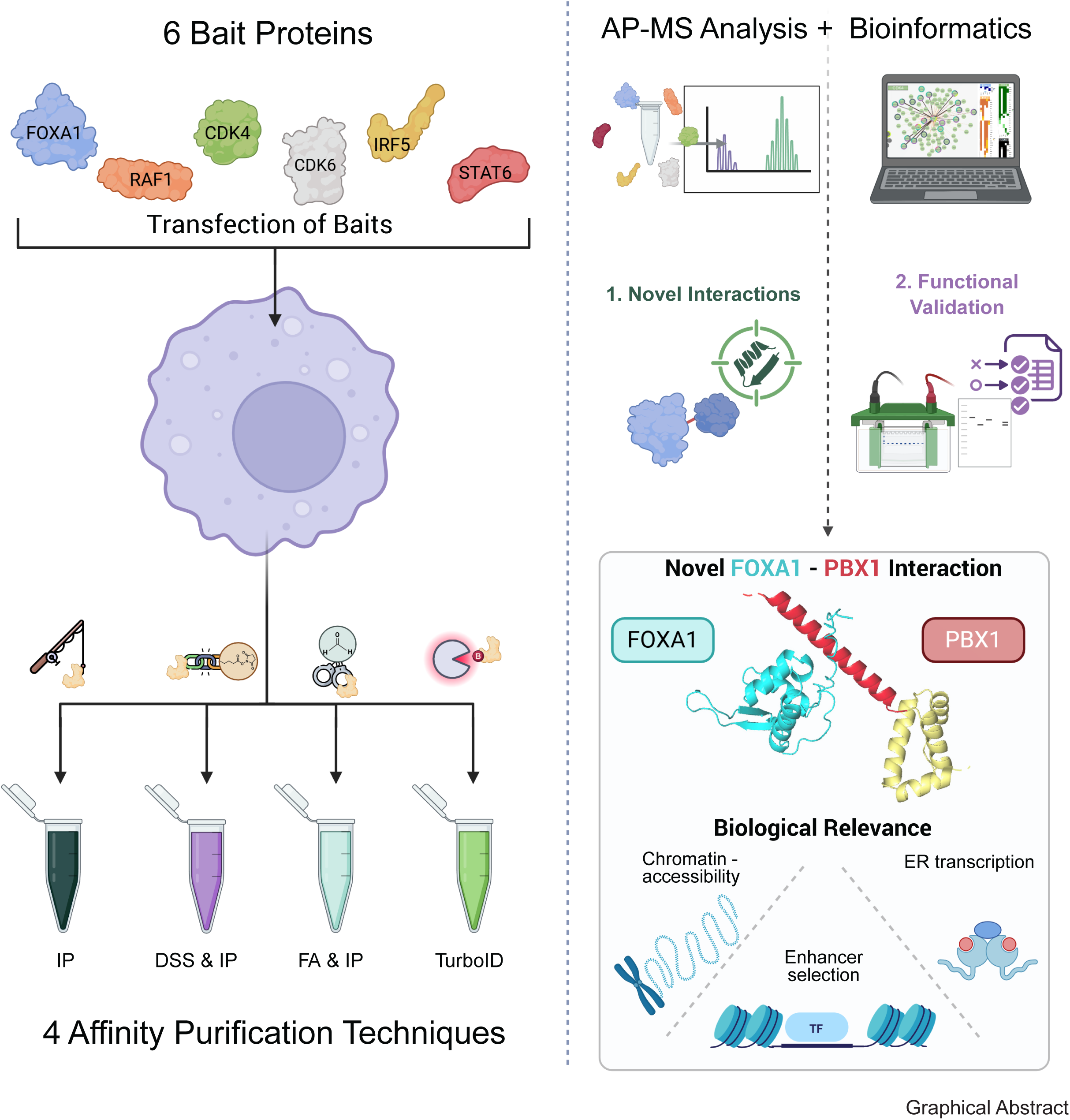

## INTRODUCTION

Protein-protein interactions (PPIs) form the structural and functional backbone of cellular networks. In cancer, aberrant PPIs underpin many hallmarks of disease, including sustained proliferative signaling, evasion of growth suppressors, altered transcriptional programs, and immune dysregulation ^1–4^. While many oncogenic drivers are well characterized at the gene or protein level, their functional impact is mediated through dynamic protein complexes whose composition, regulation, and context dependence remain often incompletely understood ^5^. Systematic mapping of PPIs therefore represents a critical step toward elucidating cancer-relevant biology and identifying new opportunities for therapeutic intervention.

Affinity purification coupled to mass spectrometry (AP-MS) has emerged as a powerful and widely used approach for interrogating protein interaction networks at scale ^6^. Classical immunoprecipitation-based strategies are well suited for capturing stable complexes but frequently fail to detect transient and/or low-affinity interactions. To address these limitations, a range of complementary approaches has been developed, including chemical crosslinking to stabilize labile interactions ^7^ and proximity-labeling strategies that capture transient interactions ^8,9^. Each method offers distinct advantages but also introduces specific biases with respect to interaction type, abundance, and detectability. As a result, individual approaches typically provide only a partial view of the interactome.

Several studies have compared subsets of affinity purification strategies or applied alternative techniques in isolation; however, systematic integration and benchmarking of multiple complementary approaches within a unified experimental framework remain limited. In particular, it is still unclear to what extent different purification strategies recover overlapping versus method-specific interaction profiles, how reliably they recapitulate known interaction networks, and whether method-specific interactors converge on shared biological functions. Addressing these questions is essential for interpreting proteomics-derived interaction datasets and for prioritizing previously unreported PPIs for downstream functional analysis.

In this study, we focus on six cellular proteins with well-established oncological relevance representing diverse functional classes and regulatory mechanisms. FOXA1 is a pioneer transcription factor that shapes chromatin accessibility and transcriptional programs, particularly in hormone-driven malignancies ^10^. RAF1 is a central kinase in MAPK signaling and a key mediator of oncogenic growth pathways ^11^. CDK4 and CDK6 are core regulators of cell cycle progression and validated therapeutic targets in multiple cancer types ^12–15^. IRF5 ^16^ and STAT6 ^17^ are transcriptional regulators implicated in immune signaling, inflammation, and tumor-immune interactions. Together, these proteins span soluble enzymes, chromatin-associated factors, and immune regulators, providing a representative set of cancer-relevant baits for evaluating complementary interactome mapping strategies.

Here, we established a unified proteomics workflow that integrates four affinity purification approaches (i.e., conventional immunoprecipitation, two chemical crosslinking-assisted strategies, and proximity labeling) to systematically interrogate the interaction landscapes of these six oncologically relevant proteins. We found that while each method relies on distinct biochemical principles and experimental conditions, their combined application enables complementary sampling of PPIs. Using this integrated framework, we benchmarked interaction recovery against curated databases and assessed functional coherence through pathway enrichment analyses, allowing evaluation of both overlap and method-specific contributions to interactome discovery. We biochemically validated a subset of previously unreported interactions and demonstrated how detection of weak PPIs can uncover biologically meaningful regulatory relationships, exemplified by a previously unappreciated physical interaction between FOXA1 and PBX1 with functional consequences in estrogen receptor-positive (ER^+^) breast cancer cells.

Together, this study provides a systematic assessment of complementary affinity purification strategies for interactome mapping in cancer-relevant contexts and establishes an integrated framework for expanding protein interaction networks beyond what is accessible with any single approach.

## RESULTS

### Integrated affinity purification strategies reveal expanded interaction landscapes

To systematically identify PPIs, we established an integrated interactomics workflow combining four complementary affinity purification strategies: standard immunoprecipitation (IP); chemical crosslinking with either 0.05% formaldehyde (FA & IP) or 1 mM disuccinimidyl suberate (DSS & IP) followed by IP; and proximity labeling using TurboID (**Figure 1A**). Each purification was performed with at least three independent biological replicates. These approaches were applied in parallel to six bait proteins: three kinases (RAF1, CDK4, and CDK6) and three transcription factors (FOXA1, IRF5, and STAT6), which were expressed in HEK293T cells and analyzed by quantitative affinity purification coupled to mass spectrometry (AP-MS).

**Figure 1.**
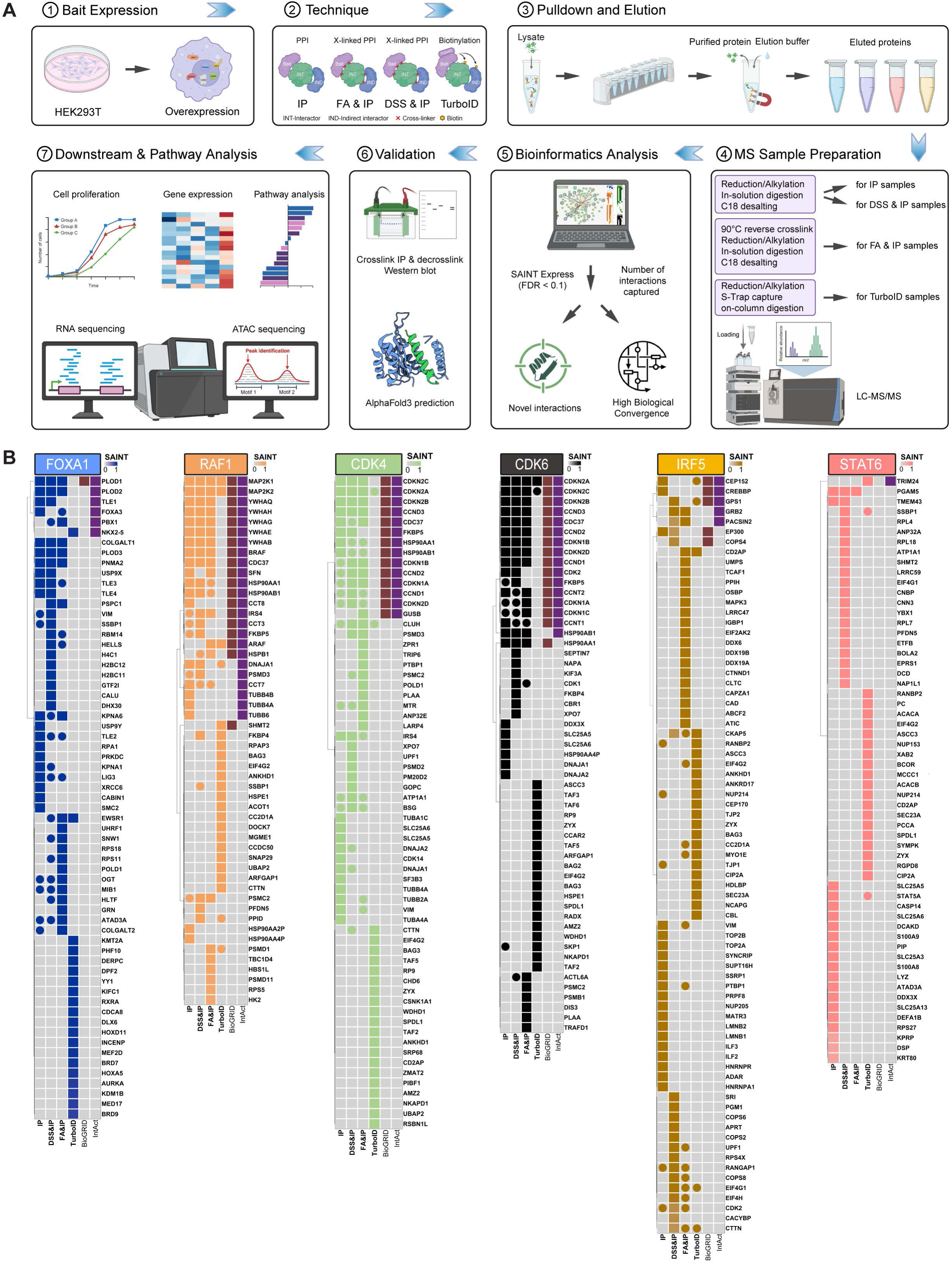
Purification workflow and top interactors. (A) Schematic overview of the experimental pipeline: (1-3) Bait proteins are expressed in HEK293T cells and processed via four complementary proteomics strategies (Standard IP, DSS/FA cross-linking, and TurboID proximity labeling). (4-5) Eluted proteins undergo specialized MS preparation and LC-MS/MS analysis, with high-confidence interactors identified via SAINT Express (FDR < 0.1). (6-7) Top candidates are validated using Western blot and AlphaFold3 prediction, followed by integrated multi-omics analysis (RNA-seq, ATAC-seq) to characterize downstream biological pathways. (B) Heatmaps depicting the top 20 interaction partners identified for each bait protein across the four affinity purification methods. Color intensity represents the corresponding SAINT score for each interaction. Cells containing a dot indicate interaction partners that were also identified as significant hits (FDR < 0.1) by at least one additional purification method but did not rank within the top 20 for the method shown. Last two columns represent the known interactors from the BioGRID and IntAct databases (brown and purple respectively), while black arrows depict the selected weak or transient interactions later in the pipeline.

This multi-strategy design was intended to maximize interaction discovery by capturing stable, transient, and proximity-based associations that are differentially accessible to individual purification techniques. This workflow yielded extensive interaction datasets, substantially increasing the number of candidate interactors compared with any single method alone (**Tables S1-S4**). SAINT analysis ^18^ of the AP-MS datasets identified hundreds of high-confidence PPIs for each bait (FDR < 0.1) (**Figure S1A**). Proximity labeling with TurboID and FA & IP consistently yielded the highest numbers of detected interactions across baits, whereas standard IP and DSS & IP identified fewer proteins. After application of the SAINTexpress FDR cutoff, all baits and techniques reached a comparable order of magnitude. The only notable exception was DSS & IP for IRF5, which detected substantially fewer high-confidence interactions, highlighting the value of complementary purification strategies in interactomics studies. Consistent protein expression and purification efficiency across all affinity purification strategies and replicates were confirmed by silver staining (**Figure S1B**). Principal component analysis (PCA) further demonstrated strong reproducibility, with biological replicates clustering by bait, supporting the robustness and technical consistency of the AP-MS workflow (**Figures S2A**).

To evaluate the contributions of different purification strategies to interaction discovery, we compared the top 20 interactors identified for each bait protein across the four affinity purification approaches (**Figure 1B**). Known interactions curated in BioGRID ^19^ and IntAct ^20^ (**Table S5**) were largely recovered in the integrated datasets. The recovery-rate of known interactions from these databases (detected at least by one method) ranged from 32.6% to 61.9% across the baits (**Figure S2B** and **Table S6**). This concordance demonstrates that the multi-method workflow reliably recapitulates known PPIs while substantially expanding the detectable interaction space.

Among the top 20 high-confidence interactors (ranked by SAINT, FDR < 0.1), many known interactions were consistently identified by multiple purification strategies. RAF1, CDK4, and CDK6 showed the strongest cross-method overlap (**Figure 1B**). We also assessed the reproducibility of the known interactions that appeared among the top 20 hits for each bait. Of these known interactions, the proportion detected by at least two distinct methods reached 50% for FOXA1, 60% for RAF1, 57.1% for CDK4, 70.6% for CDK6, and 42.9% for IRF5. In contrast, STAT6 yielded a lower absolute number of recovered hits, reflecting the relatively sparse curation currently available for this bait.

When expanding the analysis to include all significant interactors (FDR < 0.1) beyond the top 20, the cross-method validation rate (interactions identified by at least two methods) substantially increased, reaching 83.3% for FOXA1, 80% for RAF1, 92.9% for CDK4, 100% for CDK6, and 71.4% for IRF5.

In addition to recovering known PPIs, a substantial fraction of top-ranked interactions was detected by only a single purification strategy (**Figure 1B**), indicating that each method contributed unique candidate interactors to the integrated dataset.

Having defined method-specific and overlapping interaction profiles, we next specifically benchmarked each purification strategy against curated interaction databases to assess biological relevance. For each bait and method, we quantified the total number of detected proteins, significantly enriched interactors (FDR < 0.1), known number of BioGRID interactions, and their overlap (**Figures 2A** and **S3**). Fisher’s exact enrichment test revealed that crosslinking-based approaches (DSS & IP and FA & IP), consistently recovered a significant fraction of known interactions across multiple baits (**Table S7**). For baits with substantial BioGRID coverage (RAF1, CDK4, and CDK6), individual strategies (Regular IP, DSS & IP, and FA & IP) effectively reproduced known PPIs. DSS & IP analysis of IRF5, despite yielding one of the lowest numbers of significant hits (**Figure S1A**), still showed significant enrichment. Regular IP showed comparable performance in several cases, whereas TurboID detected a broader background with fewer high confidence overlaps (**Figure 2B**). All identified known PPIs annotated in the BioGRID database were visualized on a heatmap in **Figure S3**.

**Figure 2.**
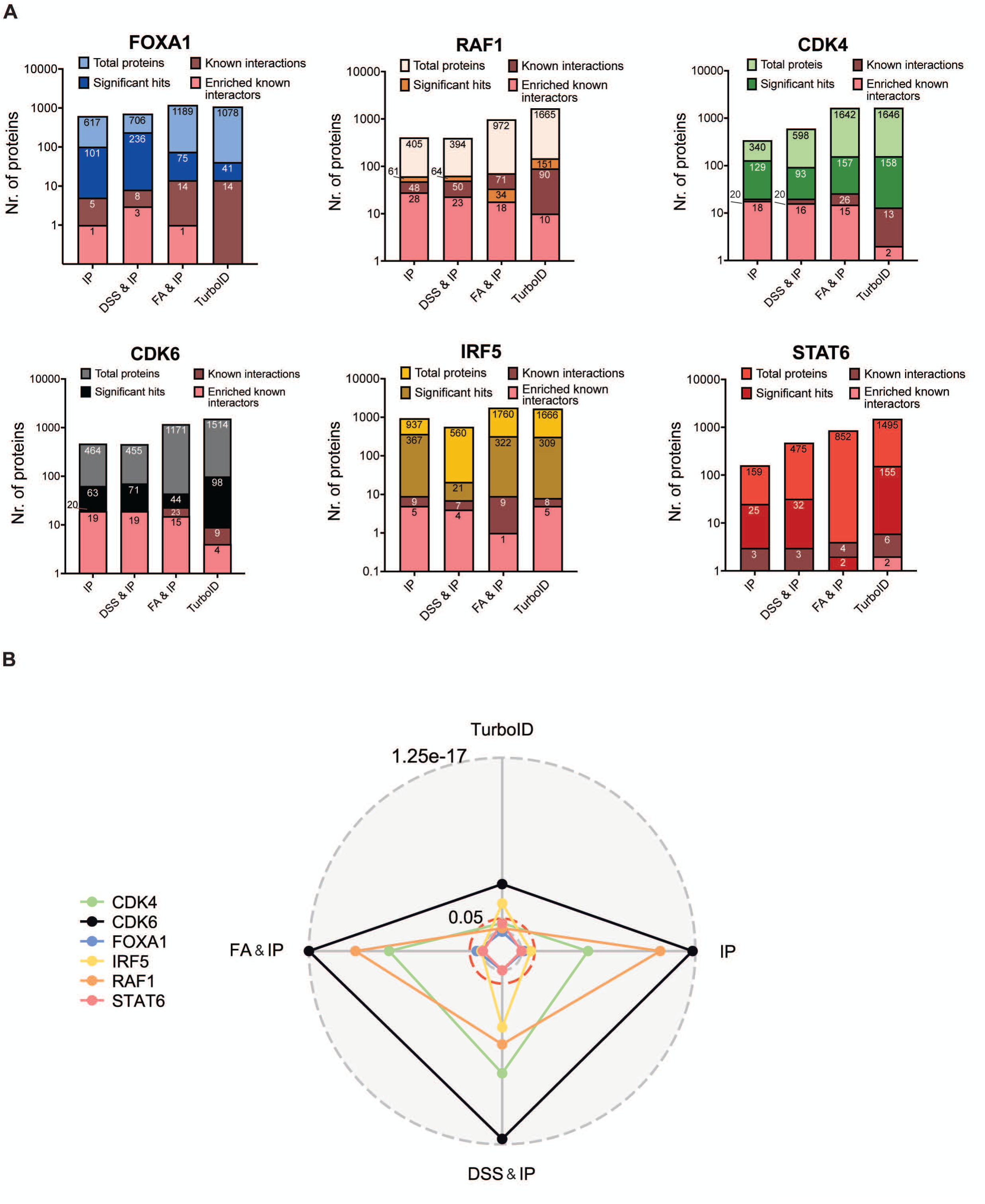
Evaluation of technique performance in recapitulating known BioGRID interactions. (A) Bar plots showing, for each bait protein, the total background of captured proteins (light bait color), significantly enriched interaction partners (dark bait color; FDR < 0.1), known BioGRID interactions present in the dataset (brown), and the subset of significantly enriched interactions overlapping with BioGRID (salmon) on a logarithmic scale. (B) Radar plot summarizing enrichment of known BioGRID interactions for each bait protein across all 24 experimental conditions. Values represent combined Fisher’s exact test enrichment scores expressed as *–log(p-values)*, reflecting the ability of each affinity purification technique to recover previously reported interactions.

To further evaluate identified PPIs using an orthogonal resource, we benchmarked our datasets against the IntAct database and compared interaction networks (**Figure S4**). First-neighbor interactors were visualized for each bait protein, and interaction confidence was assessed using IntAct MI-scores and the number of supporting experimental evidence.

Donut plot annotations integrating our results into these networks revealed that most high-confidence IntAct interactions were recovered across multiple purification strategies, while some were uniquely detected by a single method. Low-confidence edges in IntAct were correspondingly detected at lower frequency in our datasets, yet our analyses also captured additional unreported, putative weak or transient interactions.

These results are consistent with known interactions reported in both BioGRID and IntAct (**Table S5**), highlighting that our multi-method approach reliably captures previously established PPIs.

Collectively, our benchmarking analyses highlight the complementary strengths of the applied affinity purification strategies in recovering both well-characterized and unreported interaction partners. Although the protein composition of interactomes recovered by individual approaches differed substantially, functional annotation revealed strong convergence at the pathway level. KEGG and Hallmark enrichment analysis demonstrated that interactors identified across IP, DSS & IP, FA & IP, and TurboID consistently mapped to biologically coherent pathways for each bait (**Figures S5 and S6**).

### Validation of previously unreported interactions

To evaluate the robustness of the MS-based interactome dataset, we selected a panel of previously unreported candidate interactors for biochemical validation using “semi-endogenous” co-immunoprecipitation assays, in which the bait protein was exogenously expressed in HEK293T cells, immunoprecipitated and analyzed by Western blot (WB) for interaction with endogenous binding partners (**Figure 3**). Candidate selection prioritized interactions predicted to exhibit low binding affinities, as these weak and transient interactions are expected to be enriched by crosslinking methods but are often undetectable by conventional affinity purification approaches. Accordingly, we selected several interactions that were identified exclusively in the crosslinking-MS datasets, including CDK1 and HNRNPA3 for CDK4 and CDK6, PBX1 for FOXA1, FKBP4 for RAF1, GRB2 for IRF5, and LRRC59 and CLTC for STAT6. To expand the validation set and assess interactions detectable across complementary approaches, we also included candidates identified in both conventional IP and crosslinking-MS datasets, namely MTR for CDK4 and CDK6, CLUH for CDK4, HSP90AB1 for CDK6, and PGAM5 for STAT6.

**Figure 3.**
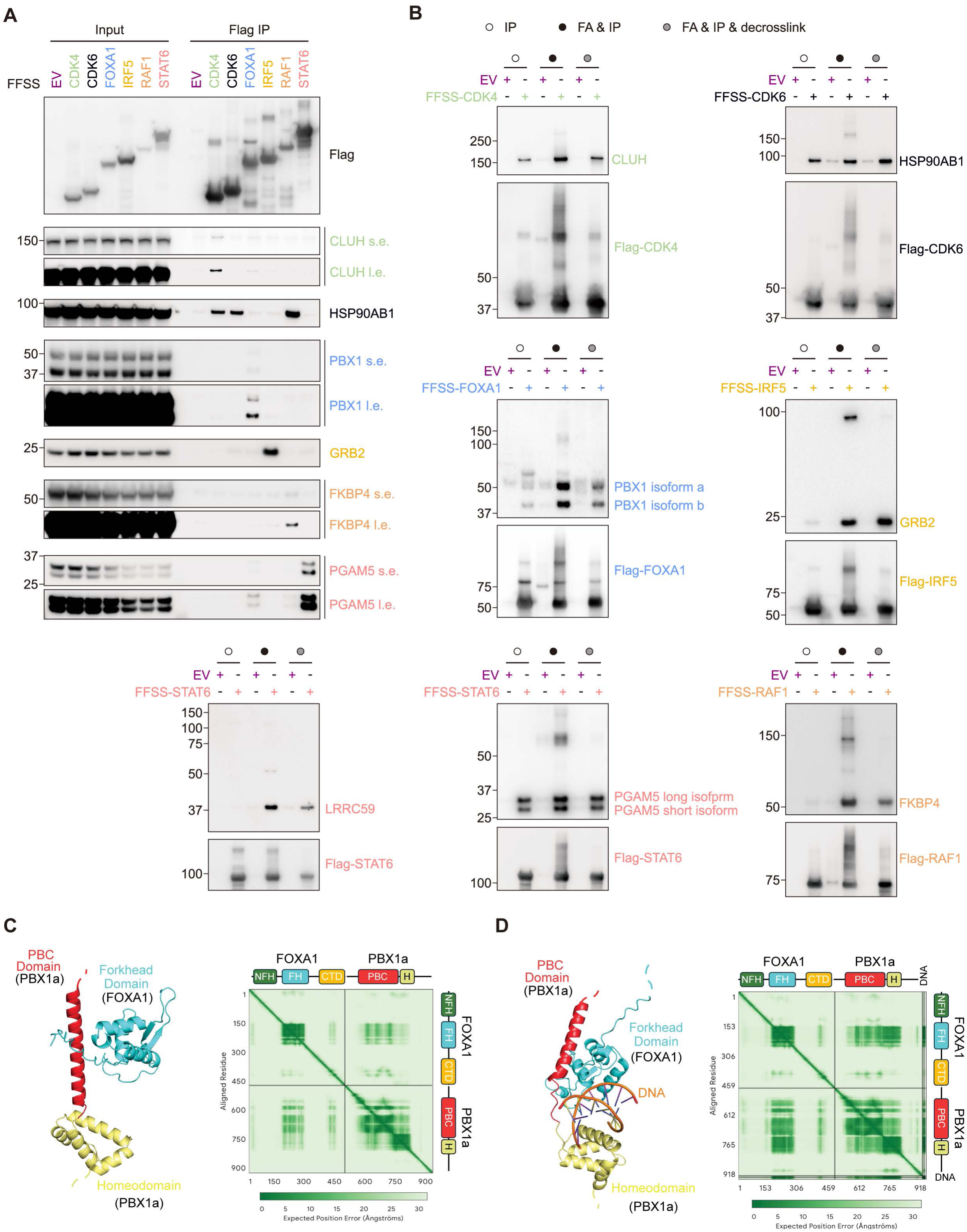
Validation of MS-identified protein-protein interactions. (A) HEK293T cells were transfected with either an empty vector (EV) or the indicated twin-FLAG- and twin-Strep-tagged proteins. Twenty-four hours after transfection, cells were harvested for immunoprecipitation (IP) and immunoblotting. S.e., short exposure; l.e., long exposure. (B) HEK293T cells were transfected with either an empty vector (EV) or the indicated twin-FLAG- and twin-Strep-tagged proteins. Twenty-four hours after transfection, cells were subjected to either standard IP or FA & IP followed by decrosslinking. (C) Left panel: Depicts the predicted interaction between the forkhead domain (cyan) of FOXA1 and the PBC (red) domain and Homeodomain (yellow) of PBX1a. Right panel: The region of low PAE at forkhead-PBC interface indicates a high positional confidence. (D) Left panel: Depicts the predicted interaction between same domains in (A) in addition to the presence of a DNA consensus sequence shared by FOXA1 and PBX1a (TGATTGAC), which enhances the positional confidence. The forkhead domain of FOXA1 interacts with the PBC domain of PBX1 and packs against the top of the dsDNA backbone. The PBC domain of PBX1 packs against the side of the dsDNA, while PBX1’s homeodomain packs against the bottom of the dsDNA backbone. Together, the three domains form interactions that wrap around the consensus sequence. The structural basis of the FOXA1-PBX1-dsDNA interaction is consistent with direct binding. Right panel: The region of low PAE is greater, indicating higher positional confidence in the FOXA1-PBX1-dsDNA interfaces.

Using this approach, co-immunoprecipitation followed by immunoblotting reproducibly confirmed 7 of the 11 selected interactions (**Figures 3A** and **S7A**), supporting the reliability of the MS-based interactome dataset. Several of these interactions were detectable by conventional IP using sensitive immunoblotting conditions and long exposure (**Figure 3A**). To evaluate whether these interactions were enhanced by crosslinking, we compared conventional IP with FA & IP either followed by decrosslinking or analyzed without decrosslinking (**Figure 3B**). Under crosslinking conditions, previously unreported interactors were captured with markedly higher efficiency than conventional IP. The STAT6-LRRC59 interaction was observed only following crosslinking. As expected, FA & IPs displayed high-molecular-weight smears, indicative of efficient crosslink formation. Following decrosslinking, proteins migrated at their expected molecular weights.

These validation experiments confirm multiple, previously unreported interactions and show that crosslinking can enhance the recovery of selected candidates that are weakly detected by conventional IP.

### FOXA1-PBX1 interaction defines a biologically relevant axis

Following biochemical validation, we asked whether the 7 newly identified interactions represent direct protein-protein contacts or are instead mediated by additional components within larger complexes. Structure-based modeling using AlphaFold3 ^21^ predicted that the FOXA1-PBX1 pair, but not the other six validated pairs, forms a direct interaction (**Figure 3C**). Notably, the confidence of the predicted interaction interface increased in the presence of DNA containing a consensus sequence recognized by both FOXA1 and PBX1 (**Figure 3D**), whereas no comparable interaction was predicted using Albumin as a negative control (**Figure S7B**). This coordinated DNA-associated binding is consistent with the established pioneer factor functions of FOXA1 and PBX1 in chromatin engagement and transcriptional regulation ^22,23^.

To investigate the functional relevance of the FOXA1-PBX1 interaction, we examined its role in transcriptional regulation in MCF7 ER^+^ breast cancer cells, where FOXA1 is a well-established pioneer factor and lineage determinant ^22,24^ and PBX1 has been shown to serve as a pioneer factor regulating ER-dependent transcription ^23,25^. While the individual roles of FOXA1 and PBX1 have been previously characterized ^22–25^, their direct physical interaction and the extent of their coordinated functions in regulating ER-dependent transcription and chromatin accessibility have not been systematically investigated. To this end, we performed targeted perturbation experiments in MCF7 cells using siRNA-mediated depletion of *FOXA1*, *PBX1*, or both factors in combination (**Figure S8A**), followed by transcriptomic analyses using RNA-seq and chromatin accessibility profiling using ATAC-seq. RNA-seq analysis identified widespread transcriptional changes across all conditions relative to the non-targeting control, with the combined *FOXA1* and *PBX1* knockdown exhibiting the greatest number of differentially expressed genes followed by *FOXA1* knockdown (**Figure 4A** and **Tables S8**). K-means clustering revealed FOXA1-dependent gene expression changes (Cluster 2) and PBX1-dependent gene expression changes (Cluster 4) with the double knockdown consistently associated with strong gene expression changes (Clusters 1 and 3) across multiple conditions, supporting an additive effect (**Figure 4B**).

**Figure 4.**
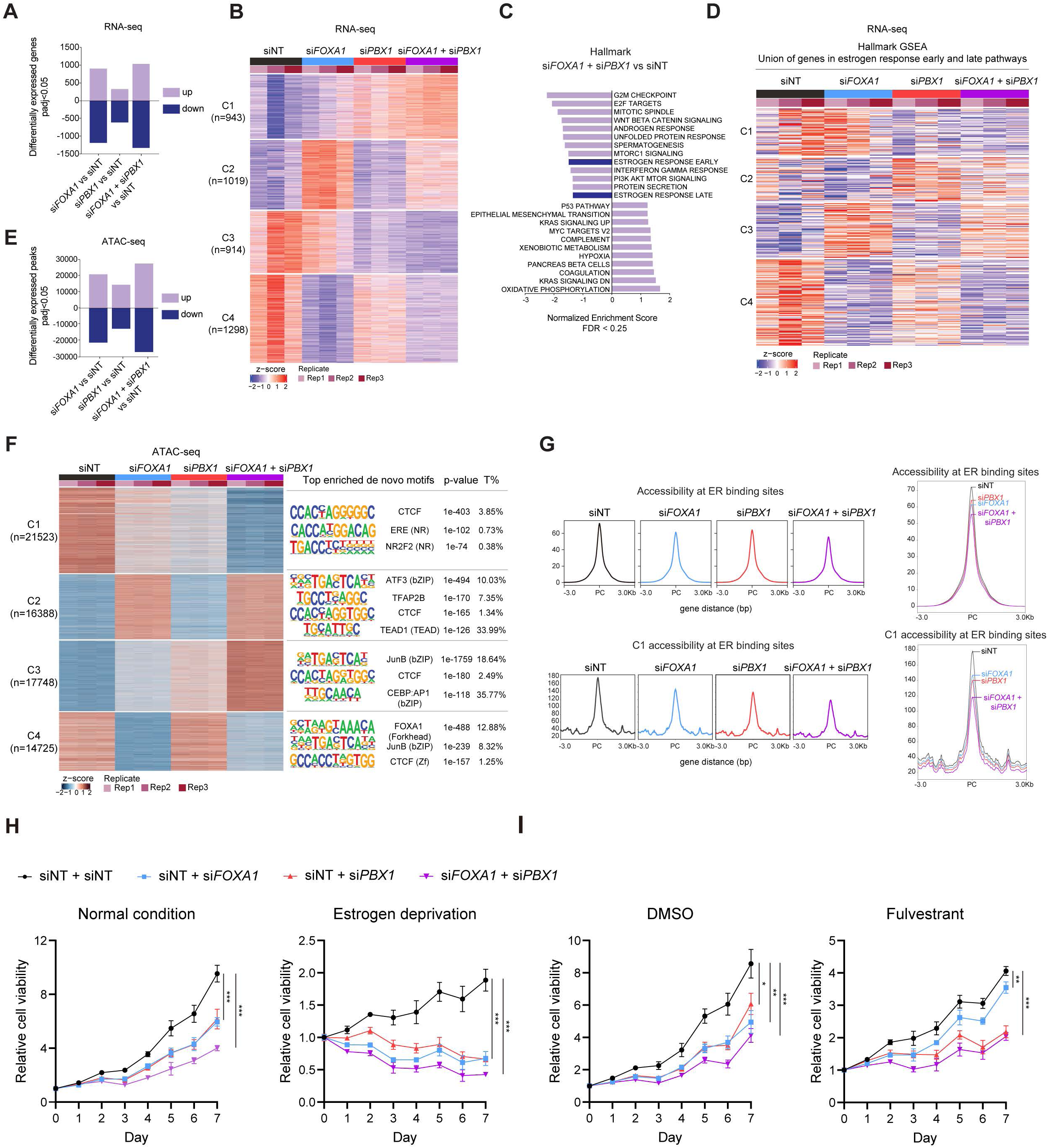
FOXA1 and PBX1 Cooperatively Regulate Estrogen-Dependent Transcription and Chromatin Accessibility in ER⁺ Breast Cancer Cells. (A) Bar plot showing the total number of differentially expressed genes (padj<0.05). (B) K-means clustering heatmap depicting gene expression changes across si*FOXA1*, si*PBX1*, and combined knockdown conditions (n = 3 biological replicates). (C) Gene set enrichment analysis (GSEA) for the comparison si*FOXA1* + si*PBX1* vs siNT. FDR < 0.25; NES, normalized enrichment score. (D) Heatmap derived from the union of genes in the Hallmark Estrogen Response Early and Late gene sets (MSigDB) across conditions (siNT, si*FOXA1*, si*PBX1*, and combination; n = 3 biological replicates). (E) Bar plot showing the total number of differentially accessible peaks (padj<0.05). (F) K-means clustering heatmap of differentially accessible peaks for each condition (si*FOXA1*, si*PBX1*, and combination; n = 3 biological replicates). Top enriched *de novo* motifs (HOMER) are shown. *p* values are indicated; %T = percentage of targets. (G) Integration of publicly available ER ChIP-seq data (GSE59530; PMID: 25752574) with accessible peaks from ATAC-seq across conditions and metaprofiles showing accessibility of cluster 1 ATAC-seq peaks at ER binding sites. Also shown is the combined metaprofiles of ER binding across the union of ER peaks overlapping ATAC-seq accessible regions. (H) Impact of *FOXA1* and *PBX1* single or combined knockdown on MCF7 cell viability in full medium or under estrogen deprivation *in vitro* (n = 6 biological replicates). Statistical significance was determined by one-way ANOVA followed by Tukey’s multiple comparisons test. Data are presented as mean ± SD. *, *P* < 0.05; **, *P* < 0.01; ***, *P* < 0.001. (I) Impact of *FOXA1* and *PBX1* single or combined knockdown on MCF7 cell viability under DMSO or 100 nM fulvestrant treatment *in vitro* (n = 6 biological replicates). Statistical significance was determined by one-way ANOVA followed by Tukey’s multiple comparisons test. Data are presented as mean ± SD. *, *P* < 0.05; **, *P* < 0.01; ***, *P* < 0.001.

Gene set enrichment analysis (GSEA) demonstrated that estrogen response pathways were among the most significantly downregulated across all perturbations (**Figures 4C** and **S8B**). Additional pathways, including EMT, oxidative phosphorylation, and MYC targets, were also affected, with *FOXA1* depletion showing the greatest overlap with the double knockdown condition. Focusing on estrogen-responsive transcription, we generated heatmaps using Hallmark estrogen response early and late gene sets, as well as the Li estrogene early response signature ^26^ (**Figures 4D and S8C**). Across these analyses, combined *FOXA1* and *PBX1* depletion resulted in the most pronounced suppression of estrogen-responsive genes.

Consistent with the RNA-seq results, ATAC-seq revealed widespread changes in chromatin accessibility across all conditions, with the most pronounced effects observed in the combined *FOXA1* and *PBX1* knockdown (**Figure 4E**), followed by *FOXA1* depletion alone (**Tables S9**). K-means clustering revealed a global reduction in chromatin accessibility, with distinct subsets of regulatory elements largely dependent on FOXA1 (Clusters 2 and 4) or cooperatively regulated by FOXA1 and PBX1 (Clusters 1 and 3) (**Figure 4F**). Notably, regions exhibiting the greatest loss of accessibility in the double knockdown compared to either single perturbation (Cluster 1) enriched for estrogen response element (ERE) motifs, whereas regions gaining accessibility were enriched for AP-1 family motifs, including JUNB (**Figure 4F**). Consistent with this, chromatin accessibility at ER binding sites, as well as sites within Cluster 1, was markedly reduced following *FOXA1* and *PBX1* depletion, with the strongest effects observed under dual knockdown conditions (**Figure 4G**).

We next performed an integrative analysis of RNA-seq and ATAC-seq data to examine the top 50 genes showing the greatest concordant changes in chromatin accessibility and gene expression, including those most strongly upregulated (**Figures S8D-S8F**) and downregulated by the knockdown of *FOXA1*, *PBX1* or the combination (**Figures S8G-S8I**). Genes exhibiting the greatest decreases in chromatin accessibility were also concordantly transcriptionally repressed, including *ESR1*, coding the ER, canonical ER targets such as *IGFBP4* and *XBP1*, consistent with their previously reported downregulation in expression studies ^25^, as well as epithelial lineage markers like *KRT8*, and *FOXP1*, a regulator of mammary ductal morphogenesis ^27^, with these effects most pronounced following *FOXA1* depletion and combined *FOXA1*-*PBX1* knockdown. Collectively, these changes support disruption of luminal identity and ER transcriptional programs. *CCND1*, a key ER target gene involved in cell proliferation, was consistently downregulated across *FOXA1*, *PBX1*, and combined perturbations. *PBX1* depletion alone, as well as the combined knockdown, also impacted broader components of the transcriptional and chromatin regulatory machinery, including *POLR2A*, *NFYC*, and *SMARCC2*.

To determine whether the observed transcriptional and chromatin-level coupling between FOXA1 and PBX1 translates into functional cellular consequences, we assessed cell viability under the same individual and combined depletion conditions. MCF7 cells subjected to *FOXA1* or *PBX1* knockdown exhibited a reduction in viability compared with control cells.

Simultaneous depletion of *FOXA1* and *PBX1* resulted in a further decrease in viability beyond that observed with either single knockdown, indicating an additive effect. Notably, this effect was maintained across normal growth conditions and endocrine stress settings, including estrogen deprivation and fulvestrant treatment (**Figures 4H** and **4I**).

Together, these findings support an additive and cooperative role for FOXA1 and PBX1 in maintaining ER-associated chromatin accessibility and transcriptional programs.

## DISCUSSION

In this study, we mapped the interaction landscapes of oncologically relevant proteins using a proteomics workflow that integrates multiple complementary affinity purification strategies. By combining conventional immunoprecipitation, crosslinking-assisted capture, and proximity labeling, we were able to sample a broad spectrum of protein associations, including stable complexes as well as putative weak, transient, or spatially constrained interactions that are often underrepresented in single-method studies. This integrated approach provides a more nuanced view of cancer-relevant protein interaction networks and highlights how methodological choice shapes interactome discovery.

Across all bait proteins, the different affinity purification strategies yielded distinct yet partially overlapping interaction profiles. Benchmarking against curated databases such as BioGRID and IntAct confirmed that known interactions were robustly recovered while simultaneously extending existing networks with previously unreported partners. Notably, many of these additional candidates were recovered by only a single purification strategy, particularly the crosslinking-assisted and proximity-labeling approaches, consistent with their ability to sample interactions that may be difficult to detect using conventional immunoprecipitation alone. Importantly, method-specific interaction profiles did not translate into divergent biological interpretations. In fact, despite some differences at the protein level, functional enrichment analyses consistently revealed convergence on coherent, cancer-relevant pathways associated with each bait. These differences likely reflect variability in interaction strength or accessibility depending on the purification strategy, underscoring the robustness of our dataset. Collectively, these findings suggest that complementary purification strategies capture different components of shared biological programs rather than unrelated interaction noise.

Targeted biochemical validation confirmed a subset of previously unreported interactions, including several that were inefficiently detected by conventional immunoprecipitation but stabilized by crosslinking. While structure-based modeling predicted direct binding for only 1 of 7 validated interaction pairs, this does not diminish the biological relevance of the remaining validated associations, many of which likely reflect protein complex-mediated interactions that cannot by captured by Alphafold3.

The FOXA1-PBX1 interaction provides a clear example of how detection of weak or previously underappreciated protein-protein interactions can reveal meaningful regulatory relationships. Structural predictions supported a direct FOXA1-PBX1 association, and functional analyses in ER⁺ breast cancer cells demonstrated that these factors cooperatively regulate chromatin accessibility, transcriptional programs, and cell fitness. FOXA1 and PBX1 have been studied in parallel or as partially overlapping chromatin regulators ^22–25^ rather than as components of the same physical regulatory complex. The present study therefore fills an important mechanistic gap by showing that FOXA1 and PBX1 are not merely co-occupying related genomic regions but can cooperate functionally. The FOXA1-PBX1 interaction is particularly notable considering prior studies establishing both proteins as pioneer regulators of ER-dependent transcriptional programs in luminal breast cancer ^22–25^. While FOXA1 and PBX1 have previously been linked through overlapping ER cistromes and enhancer-associated functions, their direct physical association had not been demonstrated.

Importantly, this finding also highlights a broader principle in chromatin biology: interactions among transcriptional and chromatin regulatory proteins are often weak, dynamic, and transient, and therefore difficult to capture using single-method affinity purification approaches, despite being functionally essential for regulatory specificity.

Our findings extend existing models of ER chromatin regulation by suggesting that FOXA1 and PBX1 can function as part of a coordinated regulatory complex rather than solely as independent pioneer factors. The additive effects of combined FOXA1 and PBX1 depletion on chromatin accessibility, estrogen-responsive transcription, and cell fitness indicate that these proteins provide both synergistic and nonredundant contributions to maintenance of the ER transcriptional state. Given the established roles of FOXA1 and PBX1 in endocrine response and metastatic progression, this interaction provides a framework for exploring how cooperative pioneer-factor assemblies shape adaptive transcriptional states in ER^+^ breast cancer.

## LIMITATIONS OF THE STUDY

This study has a few limitations. First, all proteomics experiments were performed in a single cellular context, HEK293T cells. HEK293T cells are widely used for affinity purification-mass spectrometry because of their exceptional transfectability, high protein yield, and experimental robustness; however, they do not fully recapitulate the physiological context of all bait proteins examined. For example, endogenous STAT6 is expressed at relatively low levels in HEK293T cells, which likely contributed to its comparatively weak purification performance across the tested strategies. Consequently, some interactions may be context-dependent and could differ in more physiologically relevant cell types. Second, we employed a single, streamlined extraction protocol across all affinity purification strategies to ensure comparability between methods. This choice imposes limitations for chromatin-associated proteins, including transcription factors. Although transcription factors are generally more extractable than histones, their efficient recovery can be enhanced by nuclease treatment, sonication or high-salt conditions to release DNA-bound protein complexes. However, such approaches may introduce substantial background through DNA-mediated co-purification or disrupt weak and transient protein-protein interactions, particularly in the absence of crosslinking. To balance these trade-offs, we prioritized methodological consistency over maximal recovery of DNA-bound factors. As a result, interaction coverage for chromatin-associated proteins may be underestimated, whereas soluble proteins, such as kinases, do not seem to be affected by these constraints. Third, we validated 7 of the 11 previously unreported interactions identified in this study. The lack of validation for the remaining candidates represents negative results that may have several possible explanations and does not exclude the possibility that these interactions exist but could not be detected using the tools and experimental approaches employed. Finally, although crosslinking and proximity-labeling strategies partially mitigate these limitations by stabilizing transient and chromatin-proximal interactions, no single approach can comprehensively capture all interaction types. Extending this framework to additional cell types and extraction conditions will further refine the interaction landscapes described here.

## RESOURCE AVAILABILITY

### Lead contact

Requests for further information and resources should be directed to, and will be fulfilled by, the lead contacts, Gergely Rona, and Michele Pagano.

### Materials availability

Plasmids used in the study will be deposited to Addgene and will be publicly available as of the date of publication.

### Data and code availability

All data and code needed to evaluate and reproduce the results in the paper are present in the paper and/or the Supplementary Materials. The mass spectrometry proteomics data generated in this study have been deposited to the MassIVE repository under the dataset identifier MSV000101923, and the datasets can be accessed using username (MSV000101923_reviewer) and password (Kymera2025). The data will also be available at www.proteomexchange.org under accession PXD078684. The raw RNA-seq and ATAC-seq data generated in this study have been deposited in the Gene Expression Omnibus (GEO) database under accession numbers GSE334766. All downstream analysis results are provided in the Supplementary Materials section.

## ACKNOWLEDGMENT

The authors would like to thank Robert McDonald, Richard Miller, and Nichole O’Connell of Kymera Therapeutics for their valuable feedback and insightful discussions. This work has been partially supported by NIH grant R01-CA276187 to E.T. and NIH grant R35-GM136250 to M.P. M.P. & N.Z. are investigators with the Howard Hughes Medical Institute. G.R. is supported by the Momentum Grant of the Hungarian Academy of Sciences (LP2023-15/2023), EMBO Installation Grant (IG5670-2024) and the HUN-REN Welcome Home and Foreign Researcher Recruitment Grant (KSZF-143/2023). I.S.N is supported by the 2025-2.1.2-EKÖP-KDP-2025-00007 University Research Scholarship Programme of the Ministry for Culture and Innovation from the source of the National Research, Development and Innovation Fund.

## AUTHORS CONTRIBUTIONS

G.R. and M.P. conceived and coordinated the study. Z.C., I.S.N., Q.Z., S.K., Y.G., J.O.P., J.H., and G.R. carried out the experiments. E.L. and J.E. provided technical assistance. M.A. initial SAINT analysis of proteomics. L.H. provided protocols and suggestions for crosslinking experiments. B.U., N.Z., E.T., G.R. and M.P. supervised the study. Z.C., I.S.N., G.R., and M.P. wrote the paper, with input from all authors.

## DECLARATION OF INTERESTS

E.T. reports grants and consulting fees from Astrazeneca and Menarini. M.P. is a scientific cofounder of SEED Therapeutics and an advisor for CullGen, Lumanity, Serinus Biosciences, Sibylla Biotech, and Triana Biomedicines. M.P. has financial interests in CullGen, Kymera Therapeutics, SEED Therapeutics, Thermo Fisher Scientific, and Triana Biomedicines. In the past, M.P. and N.Z. received funds from Kymera Therapeutics. N.Z. is a scientific cofounder of and has financial interests in SEED Therapeutics and Molecular Glue Labs. N.Z. serves as a member of the scientific advisory board of Synthex, Differentiated Therapeutics, and Cold Start Therapeutics with financial interests.

## METHODS

### Cell culture and transfection

HEK293T cells (ATCC) were maintained in Dulbecco’s Modified Eagle Medium (DMEM; Gibco) supplemented with GlutaMAX, 10% fetal bovine serum (FBS; Gibco), and 1% penicillin-streptomycin (Gibco) at 37°C in a humidified incubator with 5% CO_2_. MCF7 cells (ATCC) were maintained in Dulbecco’s Modified Eagle Medium/Nutrient Mixture F-12 (DMEM/F12; Gibco) supplemented with 10% fetal bovine serum (FBS; Gibco), 1% penicillin-streptomycin (Gibco), and 11.2 μg/mL insulin, human recombinant, zinc solution (Gibco) at 37 °C in a humidified incubator with 5% CO_2_. Cell lines routinely checked for mycoplasma contamination with the MycoStrip-Mycoplasma Detection Kit (Invivogen). Transient plasmid transfections were performed using Lipofectamine 3000 (Invitrogen) according to the manufacturer’s instructions, and cells were collected 24 hours post-transfection. Transient siRNA transfections were performed using Lipofectamine RNAi Max (Invitrogen) according to the manufacturer’s instructions, and cells were collected 48-72 hours post-transfection.

### Immunoprecipitation

HEK293T cells were transfected with N-terminally FFSS-tagged (2xFLAG-2xSTREP) FOXA1, RAF1, CDK4, CDK6, IRF5, or STAT6 constructs. Twenty-four hours after transfection, cells were lysed in ice-cold lysis buffer containing 50 mM Tris-HCl, pH 7.4; 150 mM NaCl; 10% glycerol; 1 mM EGTA; 1 mM EDTA; 2 mM MgCl_2_; 0.5 mM DTT; and 0.2% NP-40, supplemented with protease and phosphatase inhibitor tablets (Roche). Lysates were clarified at 16,000 × g for 10 min at 4 °C, and supernatants were incubated with anti-FLAG M2 magnetic beads (Sigma) for 2 h at 4 °C with rotation. Beads were washed twice with lysis buffer and twice with wash buffer (50 mM Tris-HCl, pH 7.4; 150 mM NaCl; 1 mM EDTA; 0.5 mM DTT; 0.2% NP-40). Bound proteins were eluted using 3×FLAG peptide (Sigma) with final concentration 100 μg/mL for 20 minutes at room temperature (RT).

### Crosslinking and affinity purification

For crosslinking, HEK293T cells were transfected with the constructs as detailed above, washed once with phosphate-buffered saline (PBS) and incubated with either 0.05% formaldehyde (FA; EMS) or 1 mM disuccinimidyl suberate (DSS; Thermo Fisher Scientific) in PBS at 37 °C for 15 min. Crosslinking was quenched with 0.125 M glycine (pH 7.0) at 37 °C for 5 min, followed by one PBS wash. Cells were then lysed in lysis buffer (50 mM Tris-HCl, pH 7.4; 150 mM NaCl; 10% glycerol; 1 mM EGTA; 1 mM EDTA; 2 mM MgCl_2_; 0.5 mM DTT; and 0.2% NP-40) supplemented with protease and phosphatase inhibitors (Roche). Lysates were cleared by centrifugation at 16,000 × g for 10 min at 4 °C. Supernatants were incubated with Strep-Tactin XT magnetic beads (IBA) for 1.5 h at 4 °C with rotation. Beads were washed twice with lysis buffer and twice with wash buffer (50 mM Tris-HCl, pH 7.4; 150 mM NaCl; 1 mM EDTA; 0.5 mM DTT; 0.2% NP-40), followed by elution with Strep-Tactin XT elution buffer (IBA) for 20 minutes at RT.

### Proximity labeling (TurboID) and purification

HEK293T cells were cultured in DMEM supplemented with GlutaMAX, 10% dialyzed fetal bovine serum (FBS; Gibco), and 1% penicillin-streptomycin (Gibco) for 3 days prior to transfection to reduce background from endogenously biotinylated proteins. Cells were then transfected with plasmids encoding either TurboID alone (control) or TurboID fused to the N-or C-terminus of FOXA1, RAF1, CDK4, CDK6, IRF5, or STAT6 using a ∼5 nm long linker (3xGGGGS). Both N- and C-terminal fusion constructs were analyzed to minimize potential steric effects of tag placement and to maximize coverage of proximal interactors. Twenty-four hours after transfection cells were incubated with 50 μM biotin for 3 h prior to collection then lysed in RIPA buffer (Thermo Fisher Scientific) supplemented with 1 mM EDTA, 1 mM DTT, and protease/phosphatase inhibitors (Roche). Lysates were cleared by centrifugation at 16,000 × g for 10 min at 4 °C, and the supernatants were adjusted to contain 250 mM NaCl. Samples were incubated with Dynabeads MyOne Streptavidin C1 (Invitrogen) for 2 h at 4 °C with gentle rotation. Beads were first washed twice with lysis buffer with protease/phosphatase inhibitors (Roche), twice with RIPA buffer containing 500 mM NaCl, and eventually twice again with RIPA buffer. A high salt buffer wash is necessary to reduce non-specific background. Bound proteins were eluted with 70 mM Tris-HCl (pH 7.4), 2% SDS, 1 mM EDTA, and 350 mM β-mercaptoethanol at 95 °C for 10 min.

### Mass spectrometry for proteomics

The samples were reduced, alkylated, and digested with trypsin. Peptides were desalted and eluted using various methods appropriate for each sample set (please refer to the supplemental section for sample prep specifics for each experiment type). The desalted samples were then reconstituted in 0.5% acetic acid and stored in the -80 until analysis.

Samples were loaded onto a trap column, connected to an EASY-Spray column using the autosampler of an Easy nLC (Thermo Fisher Scientific) and analyzed on an Orbitrap mass spectrometer (Thermo Fisher Scientific). All acquired MS2 spectra were searched against a UniProt human database using Sequest HT within Proteome Discoverer 1.4 (Thermo Fisher Scientific). The resulting peptide spectra matches and proteins were filtered to better than 1% false discovery rate (FDR) and only proteins with at least two different peptides are reported. To identify binding partners, proteins were ranked using SAINT Express. For detailed methods on mass spectrometry, see **Supplementary Methods**.

### Identification and selection of weak interaction candidates

Candidates for biochemical validation were selected through a systematic, multi-step process. First, a high-confidence candidate pool was generated for each bait, consisting of the *Top 50* interactors (decreasing SAINT & logFC, FDR < 0.1) that were identified by at least two of the four methodologies (Tables S1-S4). Our primary strategy was to prioritize previously unreported, weak, or transient interactions. This involved selecting candidates that were exclusively detected in the 0.05% FA and/or DSS cross-linking datasets but were absent from the regular IP dataset. Preference was also given to candidates demonstrating bait-specificity (i.e., unique to one bait protein). This initial list was further refined based on subcellular localization, biological relevance, and the commercial availability of validation-grade antibodies or cDNA resources. This stringent strategy successfully identified previously unreported interactions for FOXA1 (PBX1), IRF5 (GRB2), and RAF1 (FKBP4). However, due to the limited number of candidates for CDK4, CDK6, and STAT6 that met these strict criteria or passed initial validation attempts, the criteria were subsequently relaxed to also include high-confidence interactors that overlapped with the regular IP dataset. Notably, all successfully validated interactors were present in the DSS pulldown data and were independently confirmed using two orthogonal methods: semi-endogenous co-IP and a 0.05% FA cross-link/de-crosslink IP assay.

### Data visualization and enrichment analysis

For each bait, SAINT score data were filtered at FDR <0.1. Hollow heatmaps (*ComplexHeatmap* version 2.22.0) were generated to provide an overview of significant hits. Top 20 hits were selected based on decreasing SAINTScore values, and all bait proteins were discarded excluding self-interactions. Cells with dots were added to highlight hits that were significant but fell out of the top 20 cutoff for the corresponding purification method. Pathway enrichment analysis (*enrichplot* version 1.26.6; *clusterProfiler* version 4.16.4) was performed to assess functional relevance of top-ranked proteins. KEGG and MSigDB Hallmark pathway terms were plotted using the *merge* function, reporting the top five significantly enriched pathways per bait. Throughout the study, different colors were used to visually distinguish the baits: FOXA1 (blue), RAF1 (orange), CDK4 (green), CDK6 (grey/black), IRF5 (gold), and STAT6 (red).

### Benchmarking with BioGRID

Known interactors were collected from the BioGRID database (downloaded on November 12, 2024), retaining only interactions with an evidence count ≥ 2 (Table S5). No distinction was made between high- and low-throughput physical interaction types. This dataset was used to benchmark the methods applied for each bait. Heatmaps were generated to visualize overlap between known BioGRID interactors and experimentally identified proteins. For each bait, the number of interactors was quantified across five categories: the union of known interactors from BioGRID and IntAct, IntAct alone, BioGRID alone, the total proteins recovered across all four AP-MS experiments, and the subset of significant hits within those recovered proteins (Table S6). Enrichment analyses were performed using Fisher’s exact test, and radar plots were used to display *−log (p value)* results for each technique per bait (Table S7).

### IntAct network analysis

IntAct protein-protein interaction networks were accessed on August 8, 2025, and visualized in Cytoscape (version 3.10.3) ^28^ using the IntActApp plugin (version 1.1.0) ^29^. Networks contained first-neighbor interactions curated from literature, manual submissions, and IntAct data curation (Table S5). Edge weights were defined by color, indicating the MI-score (confidence metric integrating multiple factors ^30^), and by width, reflecting supporting evidence counts. For CDK4 and CDK6, which have large interaction networks, an MI-score cutoff ≥ 0.4 was applied for clarity. Networks were integrated with MS-derived results using the Omics Visualizer (version 1.3.1) ^31^.

### Hit validation using immunoblotting

Following the crosslinking and immunoprecipitation procedures, the immunoprecipitated complexes bound to beads were incubated at 65 °C overnight to reverse the crosslinks. After decrosslinking, the eluates were collected, separated under denaturing and reducing conditions on 4-12% Bis-Tris gels (NuPAGE, Thermo Fisher Scientific), and transferred onto PVDF membranes (Immobilon-P, Millipore). Membranes were blocked with 5% nonfat dry milk prepared in TBST (Tris-buffered saline containing 0.1% Tween-20) and incubated overnight at 4 °C with the indicated primary antibodies. After extensive washing with TBST, membranes were incubated for 1 h at RT with horseradish peroxidase-conjugated secondary antibodies (Amersham, GE Healthcare). Immunoreactive bands were detected using an enhanced chemiluminescence substrate (Thermo Fisher Scientific), and signals were captured using a ChemiDoc MP Imaging System (Bio-Rad).

Antibodies used in the study:

anti-CLUH (Aviva System Biology, Cat. No: ARP70642_P050, RRID: N/A)

anti-HSP90AB1 (Cell Signaling Technology, Cat. No: 4877, RRID: AB_2233307)

anti-PBX1 (Cell Signaling Technology, Cat. No: 4342, RRID: AB_2160295)

anti-GRB2 (Cell Signaling Technology, Cat. No: 3972, RRID: AB_10693935)

anti-FKBP4 (Proteintech, Cat. No: 10655-1-AP, RRID: AB_2246853)

anti-PGAM5 (Proteintech, Cat. No: 28445-1-AP, RRID: AB_2881143)

anti-LRRC59 (Bethyl Laboratories, Cat. No: A305-076A, RRID: AB_2631471)

anti-CDK1 (Cell Signaling Technology, Cat. No: 9116, RRID: AB_2074795)

anti-MTR (Proteintech, Cat. No: 25896-1-AP, RRID: AB_2880287)

anti-HNRNPA3 (Proteintech, Cat. No: 25142-1-AP, RRID: AB_2879921)

anti-CLTC (Cell Signaling Technology, Cat. No: 4796, RRID: AB_10828486)

anti-FOXA1 (Cell Signaling Technology, Cat. No: 53528, RRID: AB_2799438)

### AlphaFold driven direct interaction prediction

Structural models predicting the direct interaction between FOXA1 and PBX1 were generated using AlphaFold server with default parameters. Five ranked models were produced and assessed for structural convergence by evaluating the inter-model consistency using Cα root mean square deviation (RMSD). Model confidence was analyzed using the predicted aligned error (PAE) confidence scores (Å), which measure the relative domain and interface positioning confidence. After a direct interaction between the two transcription factors was modeled, the consensus sequences of FOXA1(5’-[AC]A[AT]T[AG]TT[GT][AG][CT]T[CT]-3’) and PBX1 (5’-TGATTGAC-3’) were assessed for a shared sequence. The consensus sequence of PBX1 is an 87% match for the FOXA1 consensus sequence parameters. Thus, 5’-TGATTGAC-3’ and its reverse complement 5’-GTCAATCA-3’ were added to a new prediction to investigate the influence of the dsDNA consensus sequence on FOXA1-PBX1 binding. The forkhead (DNA binding) domains of FOXA1 from the five solutions were superimposed, and structural variability was assessed using Cα RMSD values for the PBX1 regions contacting FOXA1. The domain and interface positioning confidence were, again, assessed using the PAE scores.

### Cell viability measurements

MCF7 cells were transfected with *FOXA1* siRNAs (L-010319-00-0005, Dharmacon) and *PBX1* siRNAs (L-019680-00-0005, Dharmacon), either individually or in combination, using RNAiMAX (Thermo Fisher Scientific). Twenty-four hours post-transfection, cells were seeded into 96-well white opaque plates with clear bottoms (Corning, tissue culture-treated) at an appropriate density. For estrogen-deprived conditions, cells were cultured in phenol red-free DMEM/F12 (Gibco) supplemented with 10% charcoal-stripped FBS, whereas control cells were maintained in complete medium. In a separate experimental set, cells were treated with either DMSO (vehicle control) or fulvestrant (Selleck Chemicals). Cell viability was assessed at the indicated time points using CellTiter-Glo (Promega). Medium was replaced for all wells on days 2 and 5.

### RNA-seq

Total RNA was isolated from frozen cell pellets using the RNeasy Mini Kit (Qiagen) with on-column DNase digestion (RNase-Free DNase Set, Qiagen), following the manufacturer’s instructions. Poly(A)-enriched human mRNA libraries were prepared and sequenced by Novogene on a NovaSeq platform (paired-end 150 bp), generating ∼6 GB of raw data per sample.

The RNA-seq data were first processed using fastp ^32^ for quality trimming of FASTQ files. Transcript quantification was then carried out with Salmon ^33^. Differential gene expression analysis was performed using DESeq2, and genes were classified as upregulated or downregulated (adjusted p-value < 0.05). Differentially expressed features between conditions were selected and subjected to k-means clustering in R following Z-score normalization. Heatmaps were generated using ComplexHeatmap ^34^, with the number of clusters manually optimized based on biological relevance. Estrogen-response gene sets from Hallmark and Li *et al* ^26^. C2 CGP were overlaid onto RNA-seq heatmaps using Entrez ID-matched genes to visualize estrogen-related expression programs across conditions. Gene Set Enrichment Analysis (GSEA) ^35^ was conducted against MSigDB pathway collections using DESeq2- normalized data (Table S8).

### ATAC-seq

Frozen cell pellets were quantified, and 50,000 cells per sample were used for transposition, following the established protocol ^36^. Libraries were generated using the Illumina Tagment DNA Enzyme and Buffer Kit (Illumina). After transposition, DNA was purified using the Zymo DNA Clean & Concentrator-5 Kit (Zymo). Libraries were PCR-amplified using 2× NEBNext Q5 Hot Start HiFi PCR Master Mix (NEB) for 11 cycles, followed by a final purification step using the Zymo DNA Clean & Concentrator-5 Kit.

For ATAC-seq analysis, raw FASTQ files were first trimmed using fastp. Reads were then aligned to the hg38 reference genome with Bowtie2 (v2.5.2) ^37^, and duplicate reads were removed using Picard’s MarkDuplicates. Peak calling was performed with MACS2 ^38^, followed by merging individual peaks to generate a unified peak atlas. Peak intensities were quantified from BAM files to produce a peak-by-sample matrix. Differential analysis, heatmap generation, and track visualization were carried out using the same workflow described for RNA-seq. Motif enrichment and peak annotation for each cluster were performed using HOMER (findMotifsGenome.pl and annotatePeaks.pl, respectively) ^39^ and profile plots were created using deepTools (v3.5.1) ^40^. External ER binding peak sets (GSE59530) ^41^ were intersected with ATAC-seq peak clusters and used as reference regions for deepTools (v3.5.1) ^40^ tornado plots and metaprofiles to compare accessibility at ER-bound sites across conditions (Table S9).

### Integration of RNA-seq and ATAC-seq

Waterfall plots were generated by integrating RNA-seq and ATAC-seq datasets to visualize the 50 most significantly up-regulated and down-regulated genes (ranked by FDR), together with their corresponding associated ATAC-seq peaks.

## Supplementary Materials

### Outline

**Supplementary Figures**

Figure S1 corresponds to Figure 1

Figure S2 corresponds to Figures 1 and 2

Figure S3 corresponds to Figure 2

Figure S4 corresponds to Figures 1 and 2

Figure S5 corresponds to Figures 1 and 2

Figure S6 corresponds to Figure 2

Figure S7 corresponds to Figure 3

Figure S8 corresponds to Figure 4

**Supplementary Methods**

## SUPPLEMENTARY FIGURES

**Figure S1.**
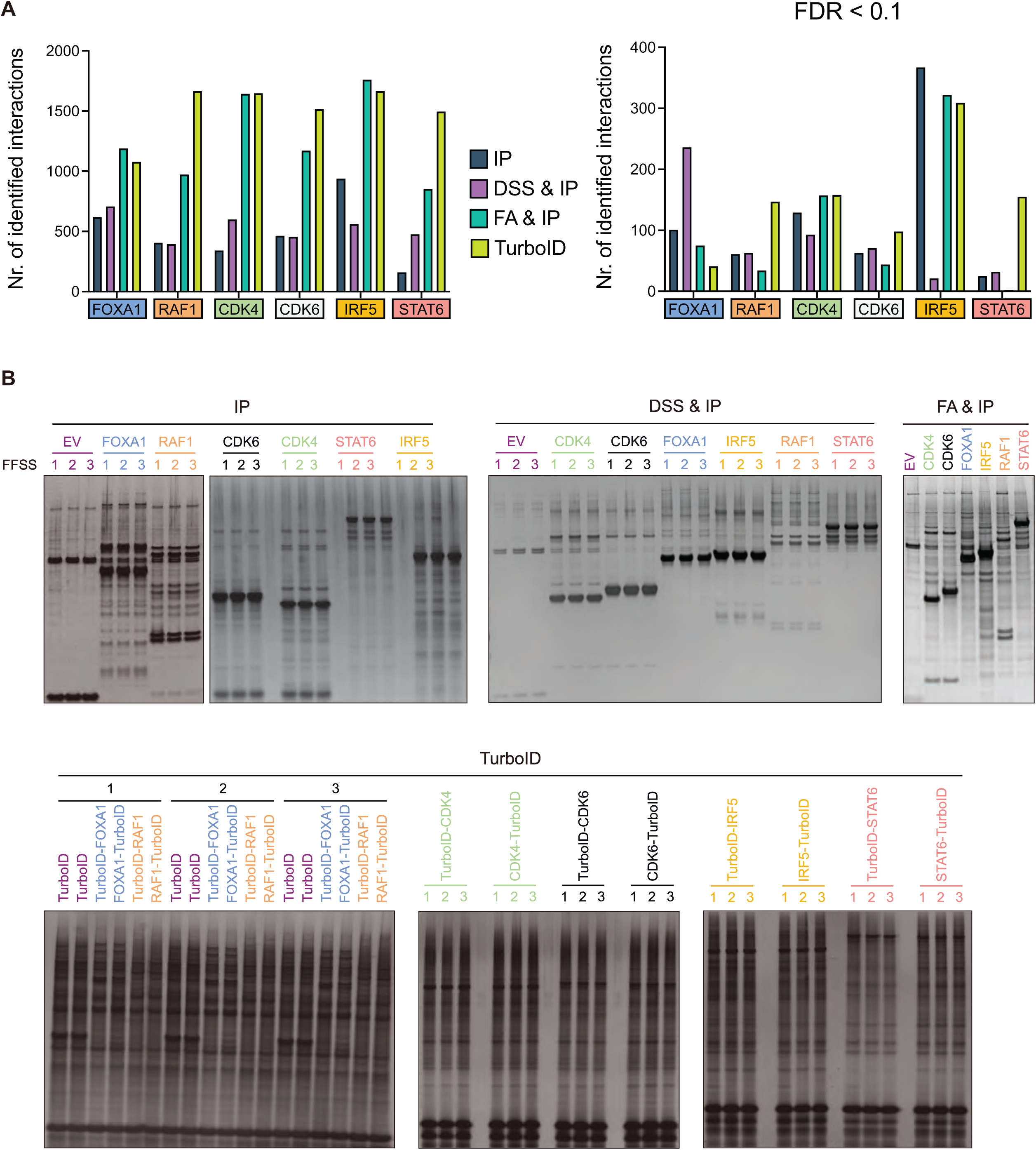
Global overview of identified, known and recovered interactions and technical validation of affinity purification approaches. (A) Bar plots summarize, for each bait protein and affinity purification technique, the total number of detected interaction partners (left) and the subset of significantly enriched interactions (right), defined by a false discovery rate (FDR) < 0.1. (B) Representative silver staining analyses demonstrating the technical performance and reproducibility of each affinity purification strategy.

**Figure S2.**
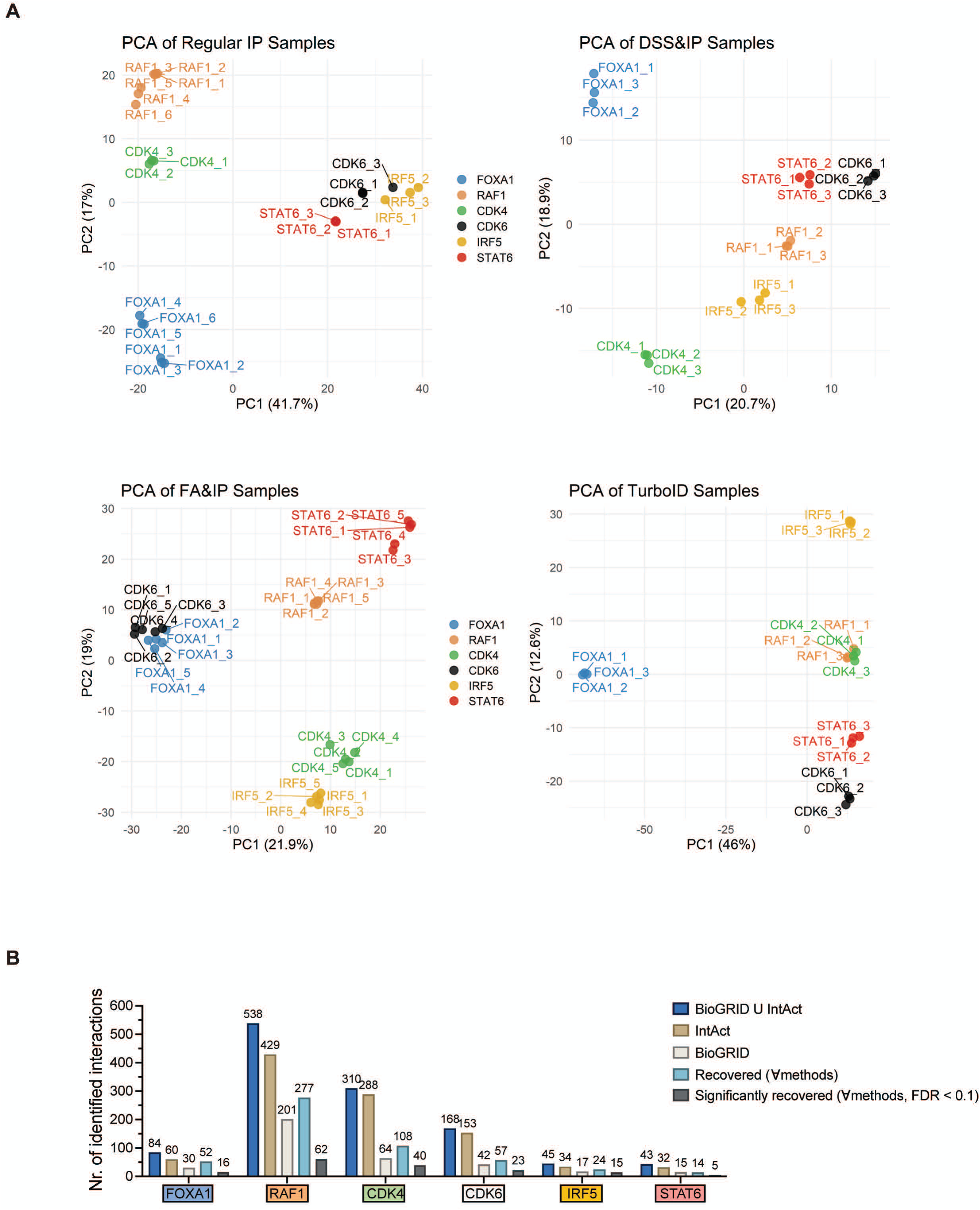
Technical replicates represent high methodological validity of each mass spectrometry run for all baits per technique. (A) Principal component analysis (PCA) plots illustrating the clustering of technical replicates from mass spectrometry analyses for all bait proteins across each affinity purification method: conventional immunoprecipitation (IP); DSS crosslinking followed by IP (DSS & IP); formaldehyde crosslinking followed by IP (FA & IP); and TurboID-based proximity labeling. Tight clustering of technical replicates demonstrates high methodological reproducibility for each technique. (B) Bar plots summarize the number of known interactors and recovered proteins for each bait. Bar heights depict the number of interactors in five categories: (1) the union of BioGRID and IntAct, (2) IntAct alone, (3) BioGRID alone, (4) the total number of interactors recovered across all four AP-MS experiments, and (5) the subset of these recovered interactors that are significant hits.

**Figure S3.**
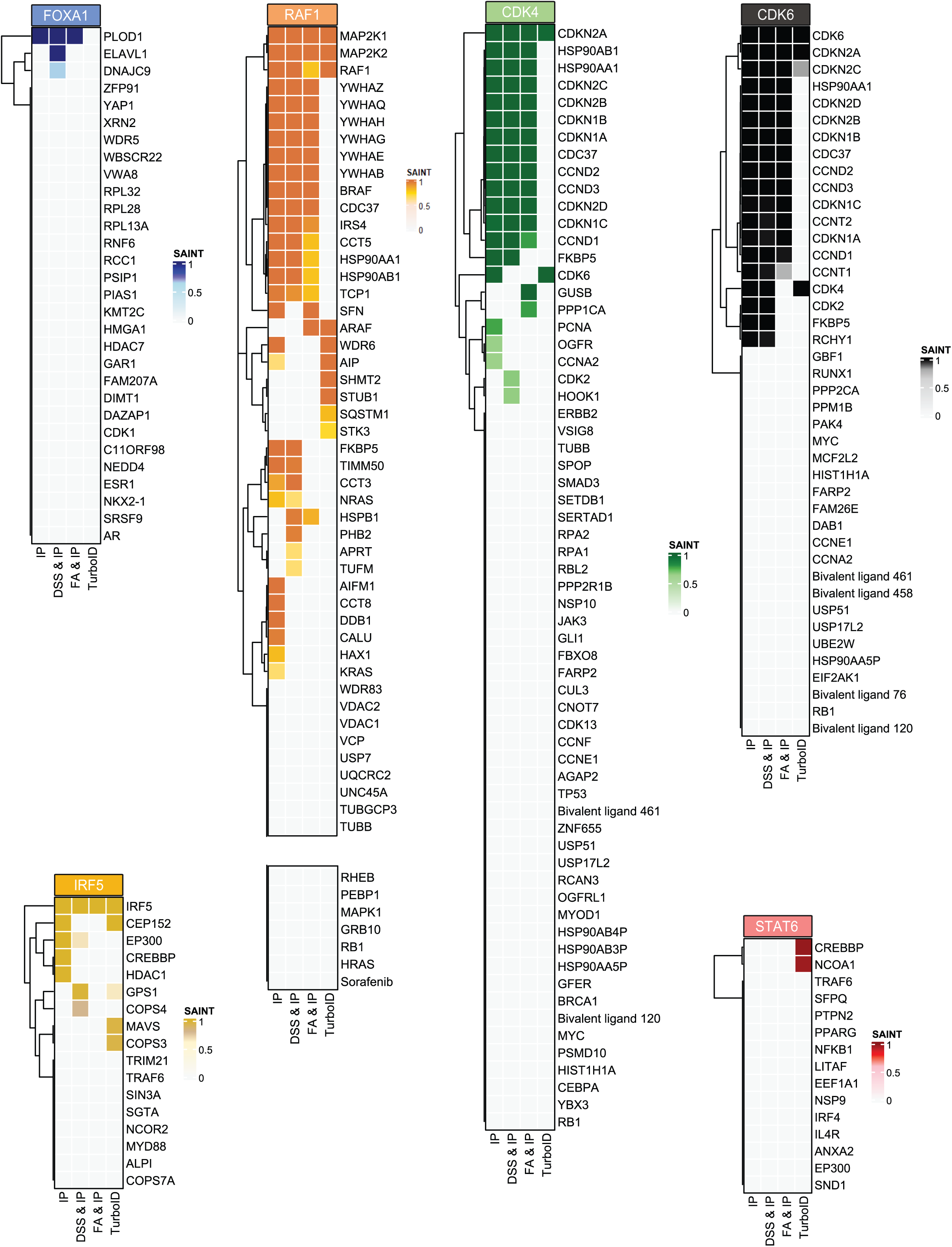
Recovery of known BioGRID interactions across affinity purification techniques. Heatmaps displaying BioGRID-curated interaction partners identified for each bait protein across all affinity purification methods. Only proteins detected by mass spectrometry in the corresponding experiments are shown, enabling direct comparison of each technique’s ability to recapitulate previously reported interactions. Color intensity represents the corresponding SAINT score for each interaction.

**Figure S4.**
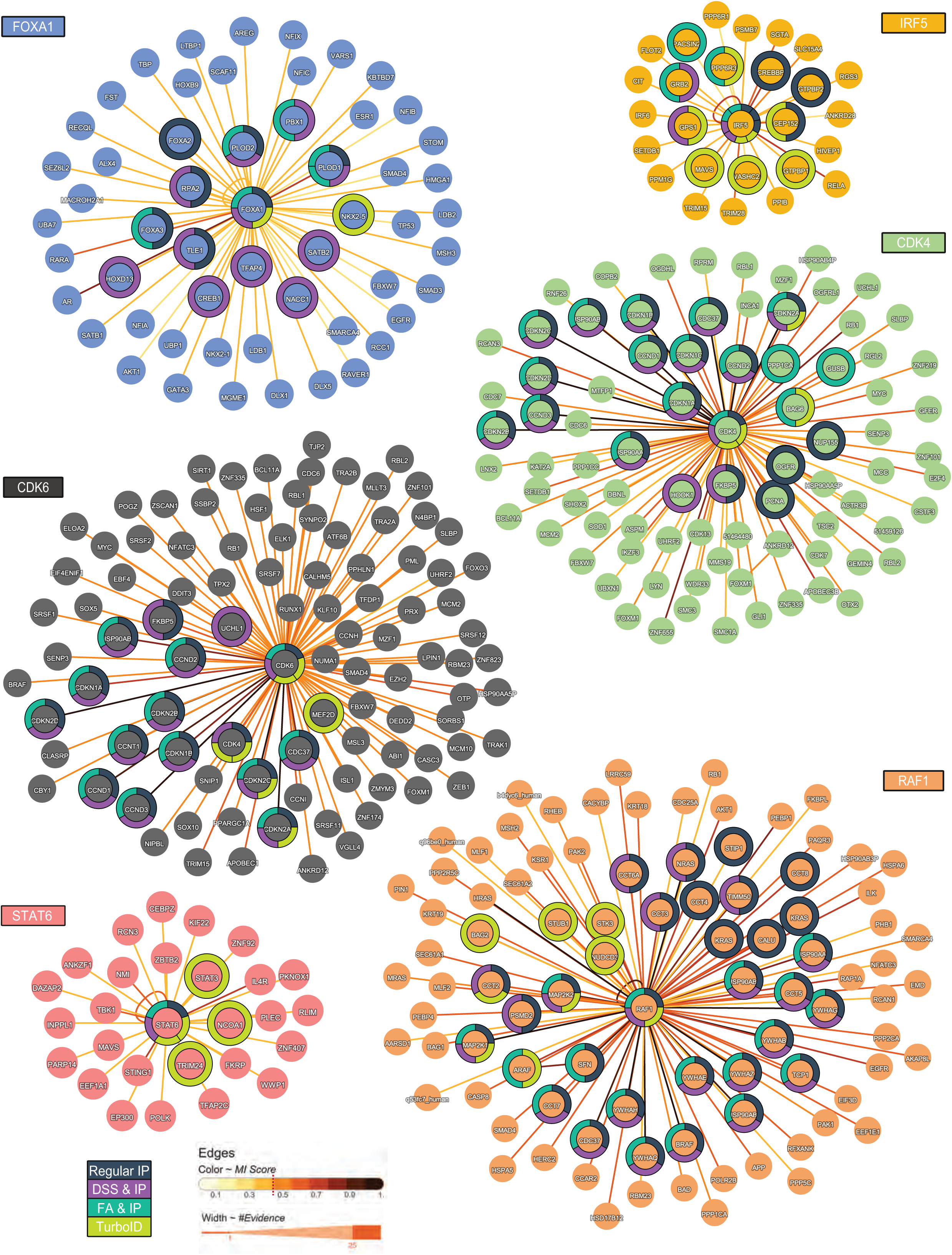
Benchmarking bait protein interactors using IntAct network analysis. Graphs show network representation of IntAct database interactors for each bait. First-neighbor interactions are visualized using IntAct’s *MI-score* and number of supporting evidence (*#Evidence*). For CDK4 and CDK6, which have large interaction networks, an MI-score cutoff ≥ 0.4 was applied for clarity. Donut coloring highlights which techniques in our study successfully captured each interactor, providing a benchmark of experimental performance.

**Figure S5.**
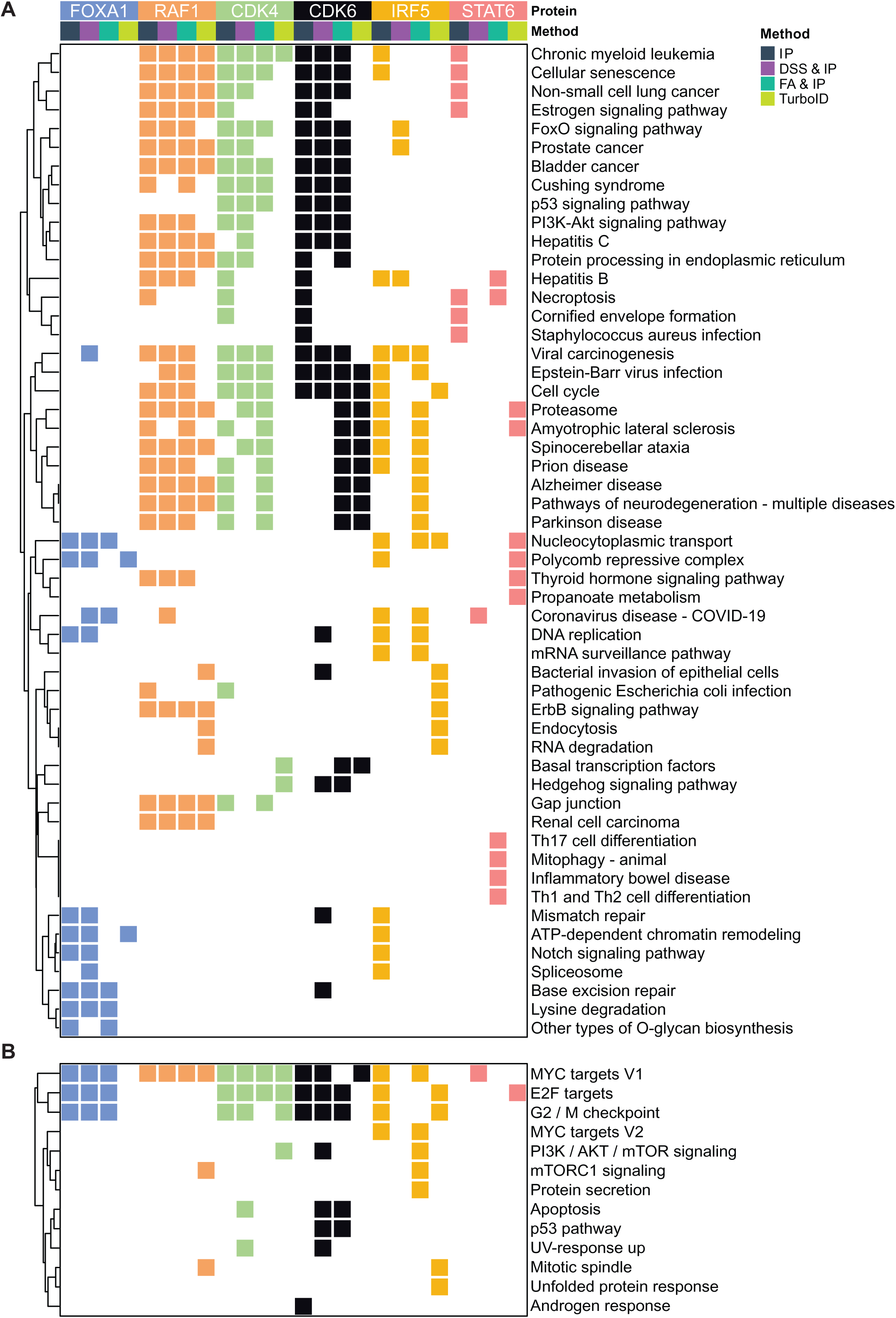
Comparative functional enrichment mapping across affinity purification techniques. (A) Heatmap showing the union of the top five enriched KEGG pathways for each bait protein identified by each affinity purification technique. Pathways ranking within the top five in any condition were included. (B) Heatmap showing the union of all significantly enriched (*adj.p* < 0.05) MSigDB Hallmark pathways for each bait protein identified by each affinity purification technique.

**Figure S6.**
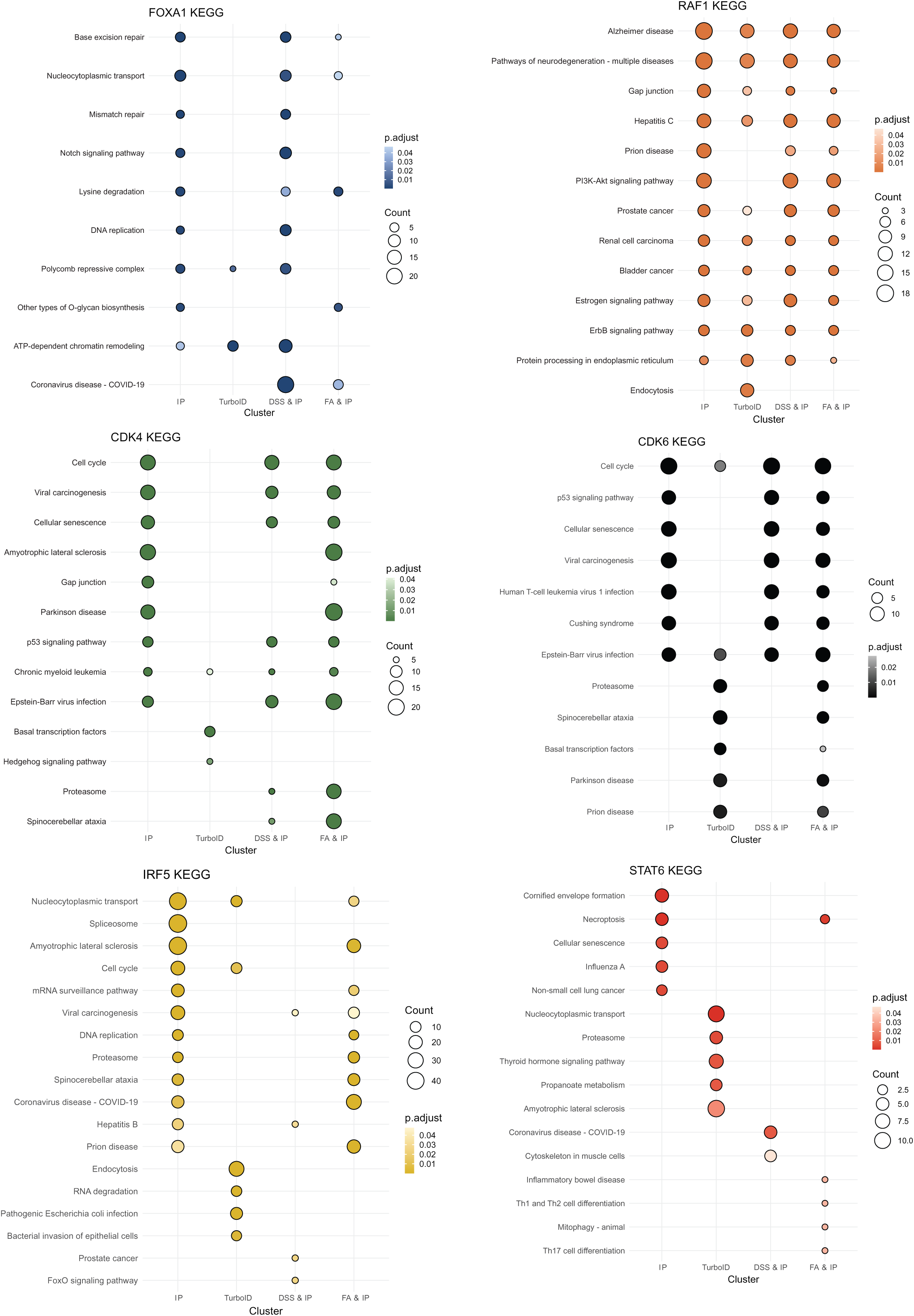
Integrated KEGG pathway enrichment across affinity purification techniques. Dot plot summarizing KEGG pathway enrichment results merged by bait protein across all affinity purification methods. For each purification technique, the top five enriched KEGG pathways were identified, and the combined non-redundant set of pathways was used for visualization. Dot color represents the adjusted p value (p.adjust), while dot size corresponds to the number of genes contributing to each enriched pathway.

**Figure S7.**
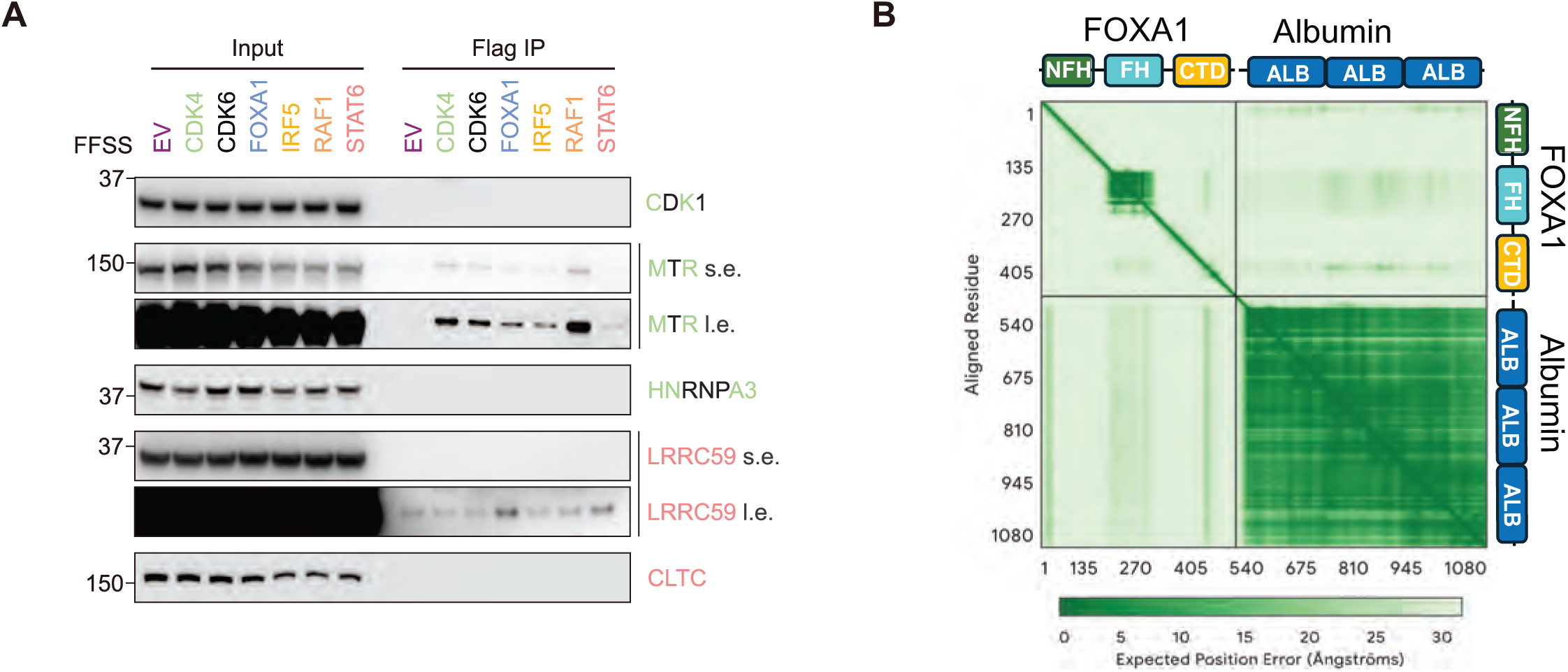
AlphaFold3 predicts FOXA1-PBX1a direct interaction: The expected position error, measured in Ångströms, depicts the confidence of an interaction. Regions of high confidence reflect interactions between structured domains. (A) HEK293T cells were transfected with either an empty vector (EV) or the indicated twin-FLAG- and twin-Strep-tagged proteins. Twenty-four hours after transfection, cells were harvested for immunoprecipitation (IP) and immunoblotting. Colors denote the corresponding bait proteins and their associated interaction hits. Alternating green and black labels indicate proteins identified as shared candidate interactors of both CDK4 and CDK6. s.e., short exposure; l.e., long exposure. (B) Depicts a nonspecific, low confidence interaction between FOXA1, a cytosolic and nuclear protein, with Albumin, a secreted protein that will not interact with FOXA1. The PAE of the FOXA1-Albumin interaction is relatively high in the forkhead and albumin domains indicating low positional confidence of the domain interfaces. The low positional confidence indicates there is no direct binding between FOXA1 and Albumin.

**Figure S8.**
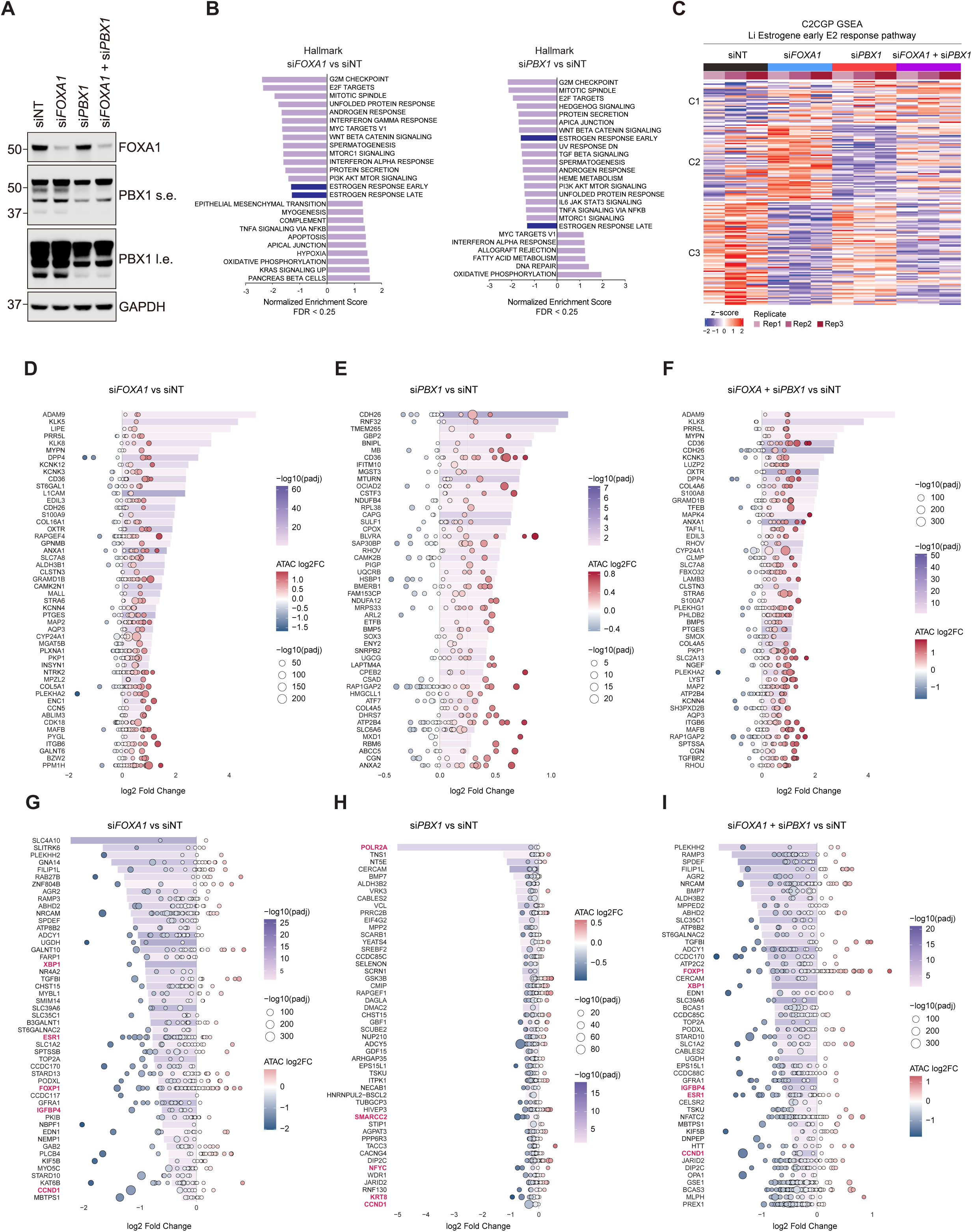
Integrated RNA-seq and ATAC-seq Analyses Reveal Concordant Transcriptional and Chromatin Accessibility Changes upon *FOXA1* and *PBX1* Depletion. (A) MCF7 cells were transfected with *FOXA1* siRNAs and *PBX1* siRNAs either individually or in combination. Forty-eight hours post-transfection, cells were harvested for immunoblotting. s.e., short exposure; l.e., long exposure. (B) Gene set enrichment analysis (GSEA) across comparisons (si*FOXA1* vs siNT, si*PBX1* vs siNT). FDR < 0.25; NES, normalized enrichment score. (C) Heatmap of genes from the C2CGP Li Estrogene Early E2 Response gene set (MSigDB) across conditions (siNT, si*FOXA1*, si*PBX1*, and combination; n = 3 biological replicates). (D-F) Waterfall plots showing the top 50 genes with increased expression associated with gains in chromatin accessibility across si*FOXA1* vs siNT (D), si*PBX1* vs siNT (E), and si*FOXA1* + si*PBX1* vs siNT (F) in full media conditions. (G-I) Waterfall plots showing the top 50 genes with decreased expression associated with loss of chromatin accessibility across si*FOXA1* vs siNT (G), si*PBX1* vs siNT (H), and si*FOXA1* + si*PBX1* vs siNT (I) in full media conditions. In the waterfall plots, the bars represent gene expression from RNA-seq datasets correlating with -log10(padj) and each bubble represents a peak whose size correlates with -log10 (padj) and color denotes log_2_ fold change.

## SUPPLEMENTARY METHODS

### Sample Preparation for Mass Spectrometry

#### Regular IP

The eluted protein samples were reduced using 2.5 μL of 0.2 M DTT for 1 h at 57 °C. The proteins were then alkylated with 2.5 μL of 0.5 M iodoacetamide (Sigma) for 45 min at RT in the dark. The proteins were then digested using 500ng of sequencing grade modified trypsin (Promega) with shaking at room temperature. Peptides were then loaded onto an equilibrated C18 Spin Column (Thermo Fisher Scientific). Peptides were washed three times with 0.1% TFA and spun on the centrifuge. Subsequent washes were done using 0.5% acetic acid and the peptides were then eluted three times using 80% acetonitrile in 0.5% acetic acid. The organic solvent was removed using a SpeedVac concentrator and the sample reconstituted in 0.5% acetic acid.

#### TurboID

The eluted protein samples were reduced using 2 μL of 0.2 M DTT for 1 h at 57 °C. The proteins were then alkylated with 2 μL of 0.5 M iodoacetamide (Sigma) for 45 min at RT in the dark. Samples were acidified with 12% phosphoric acid to 1.2%. Samples were then diluted with the S-Trap binding buffer containing (90% aqueous methanol containing a final concentration of 100 mM TEAB) and loaded onto the S-Trap (Protifi) which is placed in a 2 mL Eppendorf tube. The samples were then spun at 4,000 g for 1 min. Next, samples were then subsequently washed 3X using the S-Trap binding buffer with the spin step repeated after each wash addition. The S-trap column was then transferred to a new 1.5 mL Eppendorf tube and the proteins were then digested with 1 μg Trypsin (Promega) at 47 °C for 1 h. Peptides from the project containing the empty vector, FOXA1 and RAF1 samples were eluted by the addition of 40% acetonitrile in 0.5% acetic acid, followed by 80% acetonitrile in 0.5% acetic acid. The Other two projects containing CDK4, CDK6, IRF5, and STAT6 samples used an updated S-Trap elution protocol where the peptides were eluted once using 50mM TEAB in water pH 8.5, once using 0.2% formic acid in water, and once using 50% acetonitrile in water. Once all elution steps were complete, the organic solvent was removed using a SpeedVac concentrator and the sample reconstituted in 0.5% acetic acid.

#### DSS & IP

The eluted protein samples were reduced using 2.5 μL of 0.2 M DTT for 1 h at 57 °C. The proteins were then alkylated with 2.5 μL of 0.5 M iodoacetamide (Sigma) for 45 min at RT in the dark. The proteins were then digested using 500 ng of sequencing grade modified trypsin (Promega) with shaking at RT. Peptides were then acidified to 0.5% TFA using 10% TFA and loaded onto an equilibrated C18 Spin Column (Thermo Fisher Scientific). Peptides were washed three times with 0.1% TFA and spun on the centrifuge. Subsequent washes were done using 0.5% acetic acid and the peptides were then eluted three times using 80% acetonitrile in 0.5% acetic acid. The organic solvent was removed using a SpeedVac concentrator and the sample reconstituted in 0.5% acetic acid.

#### FA & IP

The samples were incubated at 90 °C for 10 min to remove the formaldehyde. The eluted protein samples were reduced with 2.5 μL of 0.2 M DTT for 1 h at 57 °C. The proteins were then alkylated with 2.5 μL of 0.5 M iodoacetamide (Sigma) for 45 min at RT in the dark. The proteins were then digested using 500 ng of sequencing grade modified trypsin (Promega) with shaking at RT. Peptides were then loaded onto an equilibrated C18 Spin Column (Thermo Fisher Scientific). Peptides were washed three times with 0.1% TFA and spun on the centrifuge. Subsequent washes were done using 0.5% acetic acid and the peptides were then eluted three times using 80% acetonitrile in 0.5% acetic acid. The organic solvent was removed using a SpeedVac concentrator and the sample reconstituted in 0.5% acetic acid.

#### LC-MS^2^ Analysis

For every purification method (MethodGroup1: Regular IP, TurboID; MethodGroup2: DSS, 0,05% FA and 1% FA), an aliquot of each sample was loaded onto a trap column (Acclaim PepMap 100 pre-column, 75 μm × 2 cm, C18, 3 μm, 100 Å, Thermo Fisher Scientific) connected to an analytical column (EASY-Spray column, 50 m × 75 μm internal diameter, PepMap RSLC C18, 2 μm, 100 Å, Thermo Fisher Scientific) using the autosampler an Easy nLC 1000 for MethodGroup1 and an Easy nLC 1200 for MethodGroup2 (Thermo Fisher Scientific). Solvent A consisting of 2% acetonitrile in 0.5% acetic acid and solvent B consisting of 80% acetonitrile in 0.5% acetic acid. The peptide mixture was gradient eluted using the following gradient: 5% solvent B for 5 min, 5- 35% solvent B in 60 min, 35-45% solvent B in 10 min, followed by 45-100% solvent B in 10 min. The samples were acquired for MethodGroup1 on the Q-Exactive and for MethodGroup2 on the Orbitrap Eclipse mass spectrometers (Thermo Fisher Scientific) with the following parameters: full MS spectra resolution of 70,000 (MethodGroup1) and 120,000 (MethodGroup2), an AGC target of 1e6 and 4e5 and maximum ion time of 120 and 50 ms, respectively. Scan range from 400 to 1,500 m/z for both groups. The MS/MS spectra were collected using a top 20 data dependent high resolution HCD method for MethodGroup1 and HCD activation for MethodGroup2. Corresponding parameters were: resolution of 17,500 and 30,000, an AGC target of 5e4 and 2e5, maximum ion time of 120 and 200 ms, respectively. Both groups used one microscan, 2 m/z isolation window, normalized collision energy (NCE) of 27 and a dynamic exclusion of 30 s. A first fixed mass of 150 m/z was applied only for MethodGroup1. To identify binding partners, all acquired MS2 spectra were searched against a UniProt human database using Sequest HT within Proteome Discoverer 1.4 (Thermo Fisher Scientific). Fixed modifications were set on cysteine (carbamidomethyl), variable modifications on methionine, and deamidation on glutamine and asparagine. The resulting peptide spectra matches and proteins are filtered to better than 1% false discovery rate (FDR) and only proteins with at least two different peptides are reported. AP-MS data were analyzed using SAINTExpress analysis via the REPRINT web platform (CRAPome 2.0, https://reprint-apms.org/), comparing between the baits and the empty vector to determine interacting proteins. Common contaminants were marked with a “Cont_” tag and subsequently excluded from all downstream analyses.

